# A model of colour appearance based on efficient coding of natural images

**DOI:** 10.1101/2022.02.22.481414

**Authors:** Jolyon Troscianko, Daniel Osorio

**Affiliations:** Centre for Ecology & Evolution, University of Exeter; School of Life Sciences, University of Sussex

**Keywords:** vision, visual modelling, colour appearance, visual illusions, colour constancy

## Abstract

An object’s colour, brightness and pattern are all influenced by its surroundings, and a number of visual phenomena and “illusions” have been discovered that highlight these often dramatic effects. Explanations for these phenomena range from low-level neural mechanisms to high-level processes that incorporate contextual information or prior knowledge. Importantly, few of these phenomena can currently be accounted for when measuring an object’s perceived colour. Here we ask to what extent colour appearance is predicted by a model based on the principle of coding efficiency. The model assumes that the image is encoded by noisy spatio-chromatic filters at one octave separations, which are either circularly symmetrical or oriented. Each spatial band’s lower threshold is set by the contrast sensitivity function, and the dynamic range of the band is a fixed multiple of this threshold, above which the response saturates. Filter outputs are then reweighted to give equal power in each channel for natural images. We demonstrate that the model fits human behavioural performance in psychophysics experiments, and also primate retinal ganglion responses. Next we systematically test the model’s ability to qualitatively predict over 35 brightness and colour phenomena, with almost complete success. This implies that contrary to high-level processing explanations, much of colour appearance is potentially attributable to simple mechanisms evolved for efficient coding of natural images, and is a basis for modelling the vision of humans and other animals.

## Introduction

The colour and lightness of objects cannot be recovered directly from the retinal image of a scene, but depends upon neural processing by low-level spatial filters and feature detectors along with long-range and top-down mechanisms that incorporate contextual information and prior knowledge about the visual world (Wandell 1995; Brainard and Freeman 1997; Witzel et al. 2011; Bloj, Kersten, and Hurlbert 1999). Ideally, image processing achieves lightness and colour constancy – allowing us to see colour and form veridically – but inevitably it produces visual effects and illusions, which give insight into the underlying mechanisms. Thus, the surroundings of an object affect its lightness or colour in several ways. For example, assimilation and induction effects shift appearance towards that of neighbouring colours (White 1979), whereas simultaneous contrast increases the difference between an object and the surround, and in contrast induction the surround affects the contrast of a pattern (Chubb, Sperling, and Solomon 1989; Brown and MacLeod 1997). The crispening effect – where contrasts close to the background level are enhanced – encompasses all three of these phenomena (Whittle 1992; Kane and Bertalmío 2019). Related effects in colour vision include the Abney, Bezold–Brücke, Hunt, and Stevens effects, where colours, colourfulness and contrasts shift with saturation and brightness (Fairchild 2013).

Neural mechanisms have been proposed to account for some of the foregoing phenomena, for example Mach Bands can be attributed to lateral inhibition (Ratliff 1965), brightness induction to spatial filtering in the primary visual cortex (Blakeslee and McCourt 2004), and colour constancy to photoreceptor adaptation (Judd 1940; Foster 2011) or to cortical processing (Roe et al. 2012) – but these accounts are controversial, and some effects are not easily explained (Brown and MacLeod 1997; Whittle 1992; Adelson 2000). Moreover, the lack of a comprehensive account of colour appearance limits the accuracy of the models that are typically used in design, industry and research (Hunt 2005a; Fairchild 2013).

Although photoreceptor adaptation and lateral inhibition do partly account for colour constancy and simultaneous contrast effects, their primary function is probably better understood as allowing the visual system to efficiently encode images of natural scenes, which have a large dynamic range and a high degree of statistical redundancy. Coding efficiency, which allows the brain to make optimal use of limited neural bandwidth and metabolic energy, is a key principle in early visual processing (Atick and Redlich 1992; Barlow 1961; Laughlin 1981; Ruderman, Cronin, and Chiao 1998; Simoncelli and Olshausen 2001), and here we ask how a model based on this principle might account for colour appearance.

The optimal (maximum entropy) code for natural images, as specified by their spatial autocorrelation function (i.e. second-order image statistics), approximates a Fourier transform (Bossomaier and Snyder 1986; Baddeley et al. 1997), which is physiologically unrealistic. Efficient codes can however be defined for circularly symmetrical Difference of Gaussian (DoG) or oriented Gabor-function filters, which respectively resemble the receptive fields of retinal ganglion cells and the simple cells of mammalian visual cortex (Daugman 1985; Enroth-Cugell and Robson 1966; Marĉelja 1980; Simoncelli and Olshausen 2001). In an early study, Laughlin and his co-workers (Laughlin 1981; Srinivasan et al. 1982) found that the contrast response functions and the centre-surround receptive fields of fly (*Lucilia vicina*) large monopolar cell (LMC) neurons - which are directly post synaptic to the photoreceptors - produce an efficient representation of natural images for the noise present the insect’s photoreceptor responses. Specifically, synaptic amplification at the receptor to LMC synapse and lateral inhibition between receptor outputs, give a neural code that quantitatively accords with the methods of histogram equalization and predictive coding that are used by data compression algorithms (Srinivasan et al. 1982). The centre-surround receptive fields of vertebrate retinal ganglion cells are comparable to those of fly monopolar cells (Tadmor and Tolhurst 2000), while the simple cells in visual cortex generate an efficient code for natural image statistics (Field 1987; Simoncelli and Olshausen 2001).

Our aim here is not to simulate biological vision precisely, but to model efficient coding by physiologically plausible spatial filters. We describe a Spatiochromatic Bandwidth Limited (SBL) model of early vision, which uses luminance and chromatic spatial filters at octave separations to cover the detectable range of spatial frequencies (Figures 1-3). Three parameters specify the model, namely the spatial autocorrelation function (power spectrum) of natural images, noise in the retinal signal, and the channel bandwidth – or number of distinguishable response states (Figure 1; (Laughlin 1981)). The first of these parameters is given by image statistics, the second by physiological or psychophysical measurements, and the third is estimated from psychophysical data on the crispening effect (Figure 3a; (Whittle 1992)). As the model predicts colour and lightness in naturalistic images, and accounts for various visual phenomena and illusions it offers a framework for understanding neural image processing, and is a starting point for simulating colour appearance for humans and other species.

**Figure 1.**
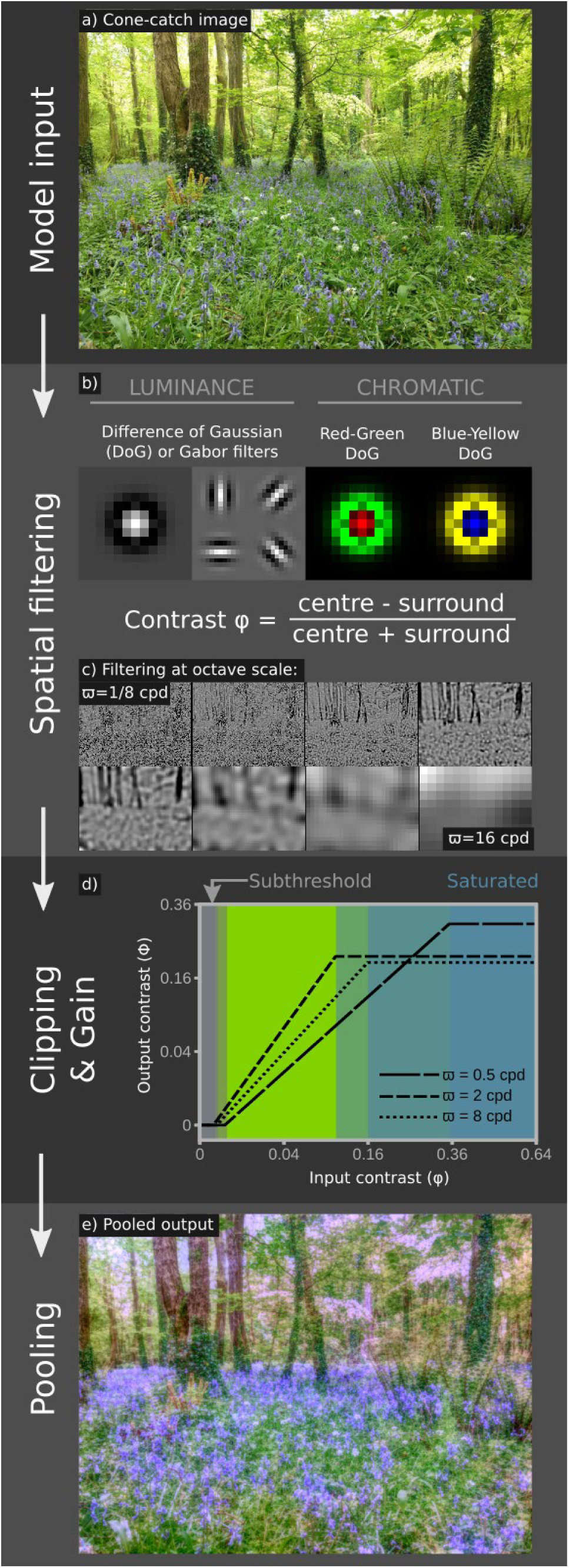
Overview of the Spatiochromatic Bandwidth Limited (SBL) model. The model uses a cone-catch image (a, Appendix), which is filtered by either DoG or Gabor kernels for luminance channels, and DoG kernels for chromatic channels (b). Contrasts are converted to Michelson contrasts (c. showing luminance DoG outputs), then clipping and gain processes are applied (d. Figure 2), and the spatial filters are pooled to create the output (e). Output colours are not scaled to sRGB space.

**Figure 2.**
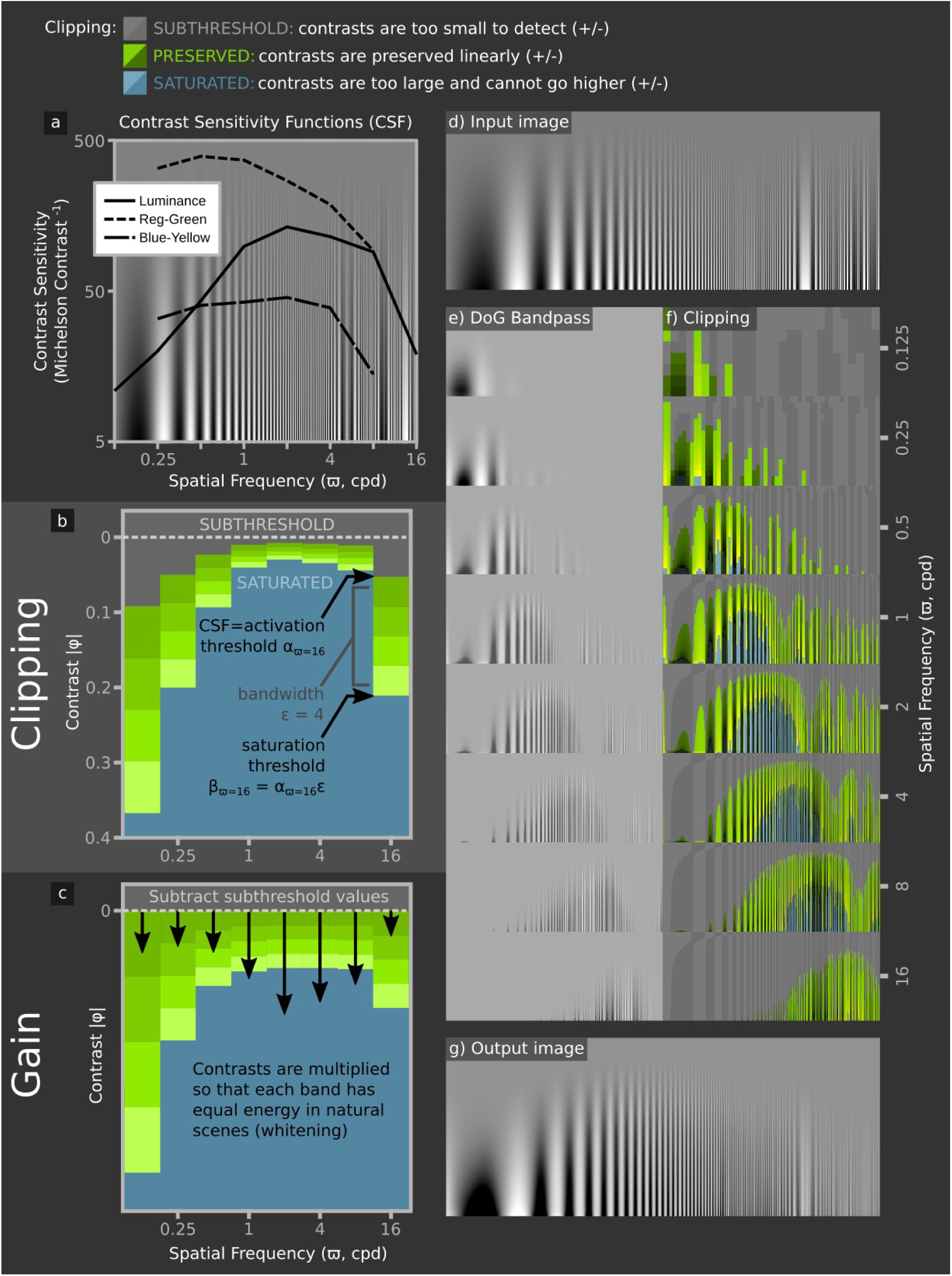
Dynamic range clipping and gain adjustment by the SBF model. a) human luminance and chromatic detection thresholds for sinewave gratings (Kim, Mantiuk, and Lee 2013). b) Clipping adjusts contrasts so that they cannot fall below the CSF at each spatial frequency (α, SUBTHRESHOLD), or above the saturation threshold (β, SATURATED). Subthreshold contrasts are subtracted, and signals at each spatial frequency are multiplied by a gain value - denoted by arrow length in (c) - so that on average natural images have equal power at each spatial frequency (whitening). The saturation threshold is calculated from the CSF and channel bandwidth, ε (4 in this example) at each spatial frequency. High and low spatial frequency channels therefore have low contrast sensitivity, but encode a large range of image contrasts, whereas intermediate spatial frequencies have high sensitivity and a low dynamic range. To demonstrate the clipping effects, we show an input image with sinewaves of different spatial frequencies and contrasts (d). (e) shows bandpass spatial filters and (f) highlights regions that are clipped or preserved. The overlap between neighbouring octaves (f) means that where contrasts are saturated for one channel, they are unlikely to be saturated for all neighbouring channels so that contrast differences are detectable even in high contrast scenes. Note that the fine lines in these illustrative images suffer from moiré effects when viewed on a monitor.

**Figure 3.**
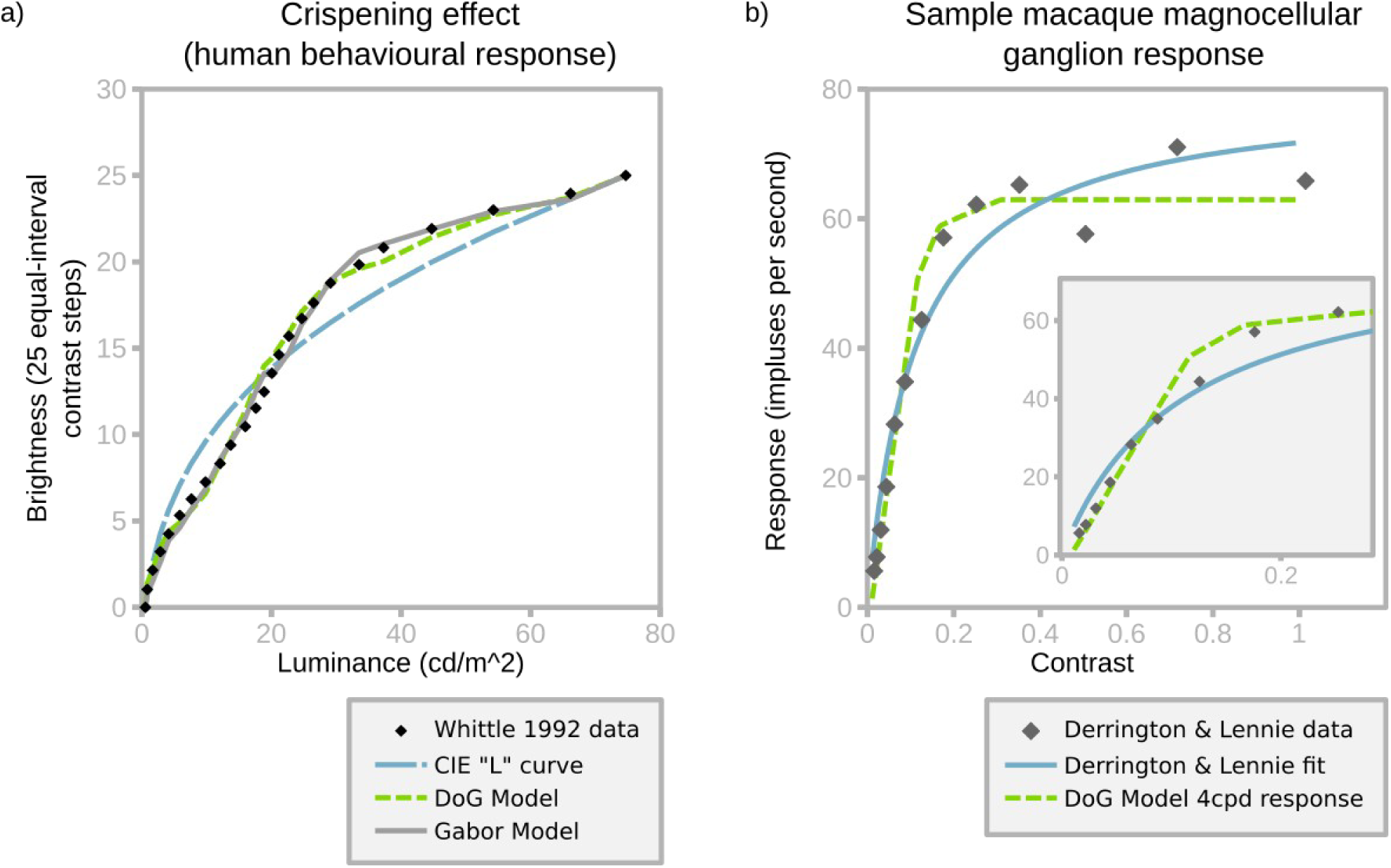
Fitting the SBL model to behavioural and neurophysiological data. a) fit to Whittle’s crispening data (1992, figure 9, “25/inc-dec/gray” treatment). Model output is scaled to the same 0 – 25 range. The best-fitting bandwidth (ε) for DoG filters is 15, and for Gabor (oriented) filters is 3.75, both of which result in a good fit to the raw data. The CIE L* fit specifies lightness in psychophysics and does not account for contrast (Commision Internationale de l’Eclairage 1978). b) Model fit to single ganglion response data from Derrington and Lennie (1989, figure 11b). Fitting used a single free parameter that multiplied the arbitrary SBL model output to match neural firing responses (with zero intercept) by least-squares regression. The SBL model shows a linear contrast response and saturation point that provide a better fit than the authors’ model. The inset excludes the three highest contrast values to highlight the linear relationship prior to saturation.

## The Model

The SBL model is comparable to other models of early vision that have been proposed to account for lightness and colour perception. These include MIRAGE (Watt and Morgan 1985), which uses non-oriented DoG filters, and the oriented difference of gaussians model (Blakeslee and McCourt 2004), which uses orientation-sensitive filters. The model differs from its predecessors in that to achieve efficient coding of natural images the gain and dynamic range (i.e. contrast response function) of neural channels vary with spatial frequency – as specified by the contrast sensitivity threshold – with gain normalised to natural scene statistics so that on average the output has equal power in each spatial channel.

The model is implemented as follows (Figures 1,2). *i*): The image is filtered with a set of spatial filters at one octave separations. These filters are either circularly symmetrical difference of Gaussian (DoG) functions (Enroth-Cugell and Robson 1966) or Gabor functions at four orientations (Daugman 1985). The filtering process differs from convolution in that it applies a Michelson contrast to centre versus surrounds. The three spectral classes of filter correspond to those in human vision, namely achromatic/luminance with centre and surround receiving the same spectral inputs, blue – yellow, and red – green with centre and surround receiving opposite spectral inputs. *ii*): The lower threshold (α) for the filter is set by the psychophysical contrast sensitivity at the filter’s centre frequency (based on contrast sensitivity functions, [CFSs], Mullen 1985; Kim, Mantiuk, and Lee 2013). α is subtracted from image contrasts, which is consistent with human psychophysics (Kulikowski 1976). The filters’ contrast response function is linear over a limited dynamic range to an upper threshold (β), which is a fixed multiple, ε, of α. ε corresponds to the number of contrast levels that can be encoded (i.e. channel bandwidth or response states; (Laughlin 1981)); (Figures 1,2). Thus, for ε = 10, the contrast saturation threshold is 10 times the activation threshold for each filter. As ε is equal for all channels, high sensitivity filters encode a smaller range of image contrasts than low sensitivity filters (Figure 2b). We estimated ε by fitting the model to Whittle’s (1992) psychophysical measurement of the crispening effect (Figure 3a, 4). *iii*) Signal power in each channel is normalised to that of the filter’s response to a natural scene, thereby whitening the average spatial frequency power spectrum of the output (Carandini and Heeger 2012). *iv*) Filter outputs are summed to recover their representation of the original image, which can be compared to human perception of the image.

**Figure 4.**
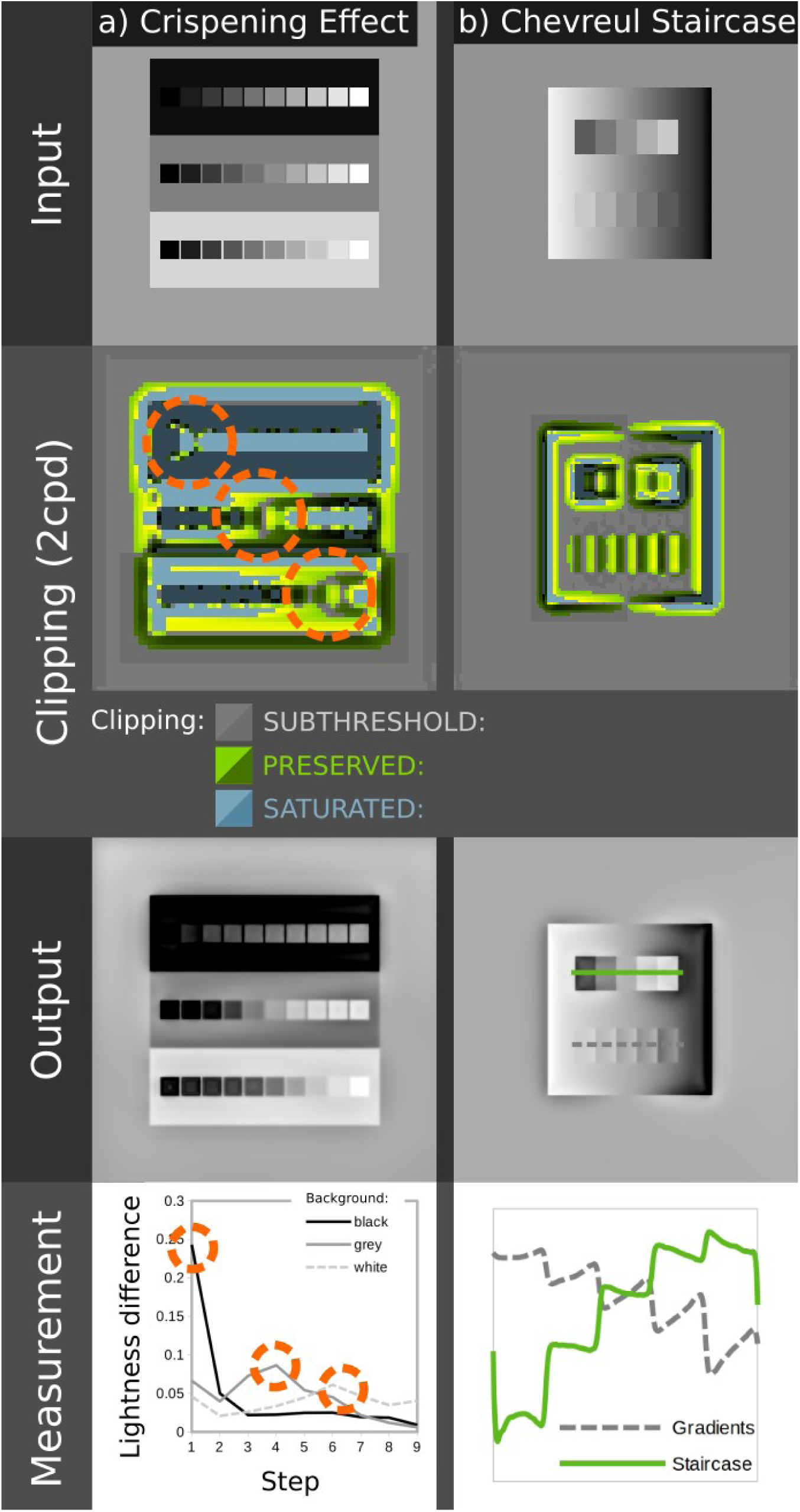
Illustration of dynamic range clipping by the SBL model. (a) for the crispening effect (Whittle 1992). The three rows of grey levels are identical, with equal step sizes. Against the black background contrasts appear largest for darker squares, whereas the opposite is true for the white background. The SBL model explains this effect through saturation; contrasts near the grey level of the local surroundings are preserved (highlighted with circles), while other contrasts are saturated (blue areas adjacent to the highlighted areas). The graph at the bottom plots differences between squares in the three rows, showing higher contrasts for dark, middle and light ranges respectively. Illusions such as the Chevreul staircase (b) are also explained in part by clipping. The upper staircase appears to be a series of square steps in grey level. The lower staircase has the same grey levels, but is flipped so that its gradient matches the surround gradient. The SBL model correctly predicts that the upper staircase is seen as square steps in grey level (solid green line) while the lower staircase is a series of gradients (dashed grey line). The plot shows pixel values in arbitrary units measured along each staircase, as highlighted in the output image. The model shows that this effect arises partly because the matched gradients of the lower staircase causes local subthreshold contrasts, and because contrasts are not balanced on each side of the step.

For the red-green and blue-yellow chromatic channels, we make the assumption, consistent with neurophysiology (Solomon and Lennie 2007; Conway 2001), that the filters are less orientation selective than for luminance channels and use only DoG filters (but see Shapley, Nunez, and Gordon 2019). The bandwidth of the red-green channel equals that of the luminance DoG signal, which produces plausible results (Figure 1 and below). However, if the blue-yellow channel has the same bandwidth (ε), its low contrast sensitivity (Figure 2a) means that it fails to saturate in natural scenes. We therefore reduced ε to give an equal proportion of saturated pixels in natural images for red-green and blue-yellow channels.

An implementation of the SBL model is provided as supplementary material for use with ImageJ, a free, open-source image processing platform (Schneider, Rasband, and Eliceiri 2012) and the micaToolbox (Troscianko and Stevens 2015; Berg, Troscianko, et al. 2020).

### Estimation of the bandwidth, ε

Channel bandwidth (ε) is estimated by fitting the model to human psychophysical data from Whittle’s (Whittle 1992) investigation of the crispening effect. This study described how perceived lightness varies with luminance, and how contrast sensitivity depends on contrast and background luminance, by asking subjects to adjust target luminances to make equal-interval brightness series (Figures 3a, 4a). We created images simulating the viewing conditions in Whittle’s experiment, including the spatial arrangement and luminance of the grey patches that he used to create perceptually uniform equal-contrast steps. Raw data (Figure 3a) were extracted from figures using WebPlotDigitiser (Rohatgi 2020). Based on least squares fitting, ε is 15 for the circularly symmetrical version of the SBF model (DoG, R^2^ = 0.994), and 3.75 for the oriented version of the model (Gabor, R^2^ = 0.995). These bandwidths are within the range encoded by single neurones (Baddeley et al. 1997; Laughlin 1981). Critically, the model recreates the characteristic inflection point around the background grey value. Lowering the bandwidth, and thereby increasing the proportion of saturated channels, produces a more extreme crispening effect, which suggests that crispening is due to saturation rather than to loss of contrast sensitivity with increasing contrast between targets and the background (Figure 2), which is the usual interpretation of Fechner’s law (Whittle 1992).

Interestingly, the model with ε derived from Whittle’s (1992) crispening data accurately predicts the responses of primate retinal ganglion cells to sinewave gratings (Derrington and Lennie 1984) (Figure 3b). The model fit (R^2^ = 0.972) is better than the authors’ own function (R^2^ = 0.952). Both the psychophysical crispening effect and bottom-up neural responses suggest that at around 4 cpd the saturation threshold for the human vision and macaque retinal ganglion cells (β_ϖ=4_) is approximately 0.2.

### Model Performance

We tested the SBL model’s ability to account for approximately thirty-seven perceptual phenomena that could plausibly be explained by low-level visual mechanisms (Adelson 2000; Shapiro and Todorovic 2016; Bertalmío et al. 2020), first for the version with oriented luminance filters, and secondarily for DoG filters (chromatic filters were always non-oriented, see above). Both versions of the model qualitatively predict almost all effects and, where relevant, their controls (Table 1, Figure 4 and Appendix). The only exceptions were the DoG (non-oriented) model’s inability to predict illusory spots and bars in the Hermann grid and Poggendorff illusions, comparatively weak performance with one control for the Chevreul staircase, and the enhanced assimilation of colour created by bars in patterned chromatic backgrounds (Monnier and Shevell 2003). Nevertheless, this performance was achieved with no free parameters (Figures 1-3), and the model can be adjusted to predict all effects.

**Table 1.**
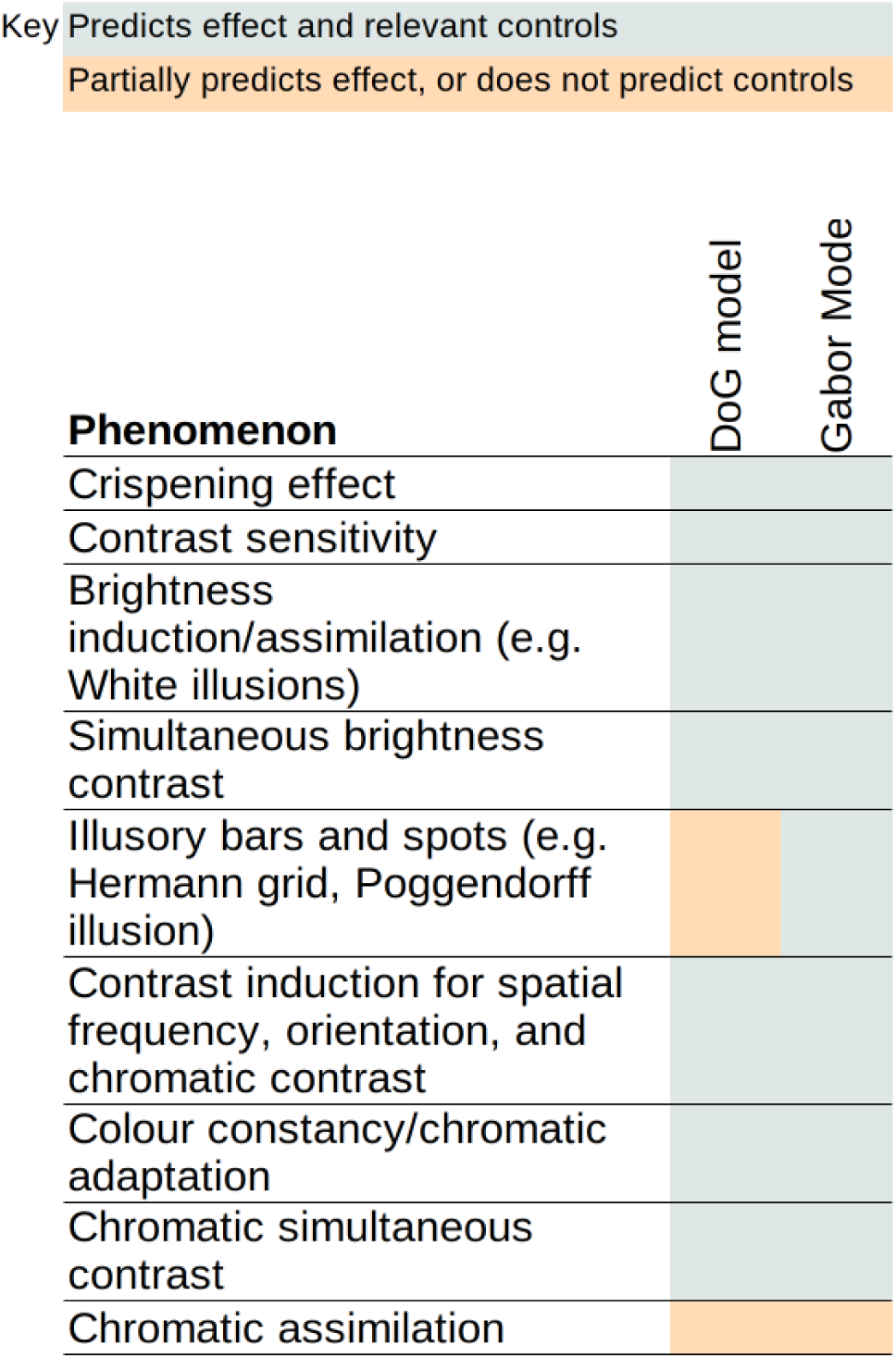
Summary of phenomena tested with oriented and non-oriented versions of the SBL model, with the parameters, α, β and ε fixed as explained in the text. All phenomena were qualitatively explained to some degree. For illustrations of specific effects see the supplementary appendix.

## Discussion

The Spatiochromatic Bandwidth Limited model of colour appearance described here at least qualitatively predicts the appearance of a wide variety of images that are used to demonstrate colour and lightness perception (Table 1, Figure 4, Appendix). These include ‘illusions’ that have been explained by high-level interpretations of 3D geometry, lighting, atmospherics, or mid-level principles of perceptual organisation (Adelson 1993; Gilchrist 2014): for example White-Munker, shadow, Koffka ring and haze illusions. It is therefore parsimonious to suggest that many aspects of object appearance can be attributed to mechanisms adapted for – or consistent with – coding efficiency (Barlow 1961). Other accounts of the same phenomena invoke specialised mechanisms (e.g. Land and McCann 1971; Blakeslee and McCourt 2004) or top-down effects, which imply that multiple sources of sensory evidence and prior knowledge are used to infer the most likely cause of the stimulus (Brown and MacLeod 1997; Gregory 1997; Yuille and Kersten 2006; Adelson 2000). Neither does the SBL model invoke light adaptation or eye movements, which implies that colour constancy is largely independent of the adaptation state of the photoreceptors – provided that they are not saturated. By comparison the models used by standard colour spaces, such as CIE LAB/CIE CAM implement the von Kries co-efficient rule (Foster 2011), which assumes that photoreceptor responses are adapted to the global mean for a scene, even though chromatic adaptation is affected by both local and global colour contrasts (Kraft and Brainard 1999). Retinex (Land and McCann 1971) and Hunt models do normalise receptor signals to their local value (Hunt 2005a) but the weightings of global and local factors are poorly understood and the underlying mechanism is unclear (Kraft and Brainard 1999). Moreover, the adjustments required for colour constancy are largely complete within about 25ms (Rinner and Gegenfurtner 2000), which is too fast for receptor adaptation, but consistent with the purely feed-forward character of the SBL model. Figure 5 shows how the SBL model can account for colour appearance in a naturalistic image under variable illumination. More generally, the feed-forward architecture of the SBL model explains why many other visual phenomena appear without any delay, whereas existing models require feedback loops for normalisation (Land and McCann 1971; Hunt 2005a; Blakeslee and McCourt 2004). Thus, Brown and MacLeod (Brown and MacLeod 1997) comment that the distribution of surround colours affects colour appearance almost immediately, leaving little time for feedback or adaptation. Likewise, as suggested by (Chubb, Sperling, and Solomon 1989), contrast induction is explained without requiring the feedback invoked by (Nassi, Lomber, and Born 2013). This is because, according to the SBL model, low contrast surrounds allow all spatial bands to operate within their dynamic ranges, whereas high contrast surrounds saturate some spatial bands, resulting in under-estimates of brightness contrast or chromaticity (Figures 3a, 4). The model also reconciles contrast constancy with a visual system that varies dramatically in contrast sensitivity and contrast gain across spatial frequencies, allowing suprathreshold contrasts to have a similar appearance at different distances (Georgeson and Sullivan 1975). Contrasts are predicted to be most constant where they are saturated across multiple spatial frequencies, e.g. where the blue regions in Figure 2f overlap. Pooling across spatial scales might explain the Abney effect, which is a shift in hue that occurs when white light is added to a monochromatic stimulus (Burns et al. 1984), because the colour stimulus may be below-threshold at some spatial bands, but above threshold for others, but we require specific data to estimate the bandwidth of chromatic channels (equivalent to Whittle’s (1992) luminance crispening data). As noted above (Model, Figures 1, 2a), we assume that the bandwidth of the red-green signal equals the luminance DoG signal, but the blue-yellow signal has reduced the bandwidth, which produces plausible results when processing natural scenes, but future work should measure the chromatic bandwidth functions and determine whether the SBL model can account for the Abney effect quantitatively. Further developments of the chromatic SBL model should also investigate whether performance could be improved by modelling both single-opponent and double-opponent pathways. The latter are sensitive to both spatial frequency and orientation, and has been suggested to play a role in suprathreshold colour appearance (Shapley, Nunez, and Gordon 2019). However, we were able to simulate the same spatial-frequency/saturation effects with the non-oriented version of the SBL model (Appendix).

**Figure 5.**
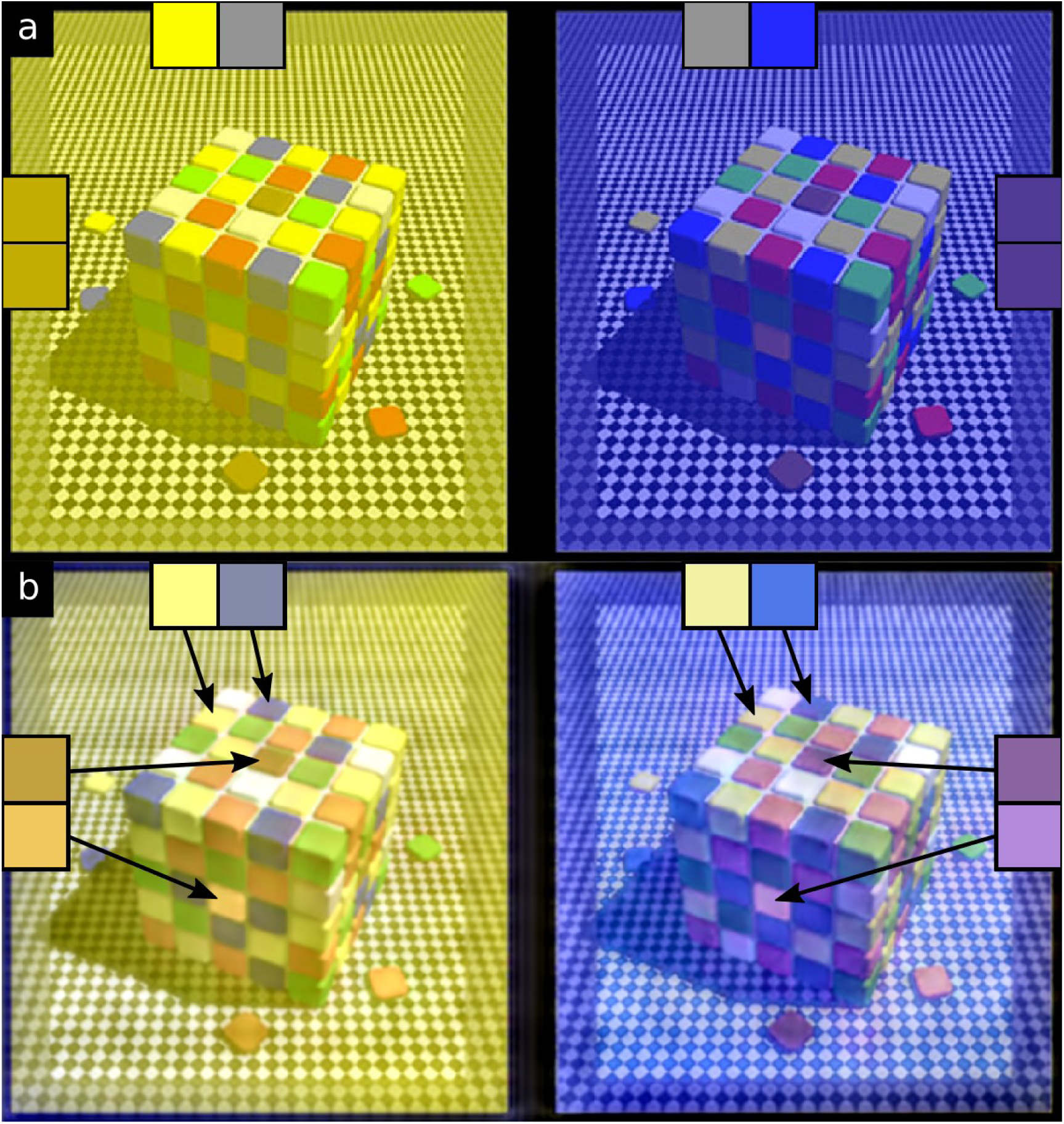
The SBL model can account for colour appearance in complex naturalistic images. (a) shows the input image (from Purves, Lotto, and Nundy 2002/Wikimedia) where the blue squares on the yellow-tinted side (left) and the yellow squares on the blue-tinted side (right) are physically the same grey (colours are shown in the squares at the top of the image). The SBL model (b) correctly predicts that the squares under both tinting regimes appear yellow and blue, rather than grey. The SBL model also predicts the powerful simultaneous contrast (or shadow) illusion present in this image whereby; the central tiles on top of the cube appear to be darker than the central tiles on the shaded side of the cube (colours shown in squares on the far left and for right hand sides).

### The Circularly Symmetric Version of the SBL Model and Animal Vision

Whereas the oriented version of the SBL model uses orientation selective achromatic filters and circularly symmetrical chromatic filters (see above), the circularly symmetrical version uses DoG filters for all channels. For the visual phenomena that we have tested the oriented version of the SBL model predicts lightness and colour at least as well as the circularly symmetrical version (table 1). It might therefore seem logical to consider only the former, but visual systems of all animals probably have circularly symmetrical receptive fields [e.g. (Srinivasan et al. 1982)], but there is limited evidence for orientation selective cells other than in mammalian visual cortex. Also, the differences between the two versions of the SBL model seem to us to be surprisingly small. For example, both predict White effects, which might be expected to depend on orientation selective mechanisms (supplementary appendix; Blakeslee and McCourt 2004; Bertalmío et al. 2020), but only the oriented model correctly predicts the presence of illusory spots in the Hermann grid, and elimination of these spots in the wavy grid (Geier et al. 2008). Similarly, the oriented version of the model predicts Koffka rings and the Chevreul staircase (Figure 4b) more accurately than the circularly symmetrical version. The bandwidth, ε, for the non-oriented filter is approximately 15, which matches neurophysiological measurements from primate retinal ganglion cells (Figure 3; Derrington and Lennie 1984). By comparison the bandwidth of the oriented version is estimated to be about four-fold lower than that of the non-oriented model, which is consistent with the low spike rates of neurons in the primary visual cortex (Baddeley et al. 1997). For a given spike rate partitioning the information into multiple channels allows a correspondingly reduced integration time.

The SBL model is useful for non-human animals because coding efficiency is a universal principle, and contrast sensitivity functions are known for many species [Figure 2a; (Caves and Johnsen 2018)], whereas psychophysical and neurophysiological data on visual mechanisms in non-primates is limited. Current research into non-human colour appearance typically uses the receptor noise limited (RNL) model (Vorobyev and Osorio 1998; Renoult, Kelber, and Schaefer 2017), which also assumes that early vision is constrained by low level noise. Others have sought to control for acuity and distance dependent effects (Caves and Johnsen 2018; Berg, Troscianko, et al. 2020; Barnett et al. 2018), but surprisingly few studies have utilised contrast sensitivity functions (Melin et al. 2016), and behavioural validation of the models is difficult (Silvasti, Valkonen, and Nokelainen 2021; Berg, Hollenkamp, et al. 2020). As with human vision, the SBL model may reconcile a number of key effects. For example, in a bird (blue tit, *Cyanistes caeruleus*) chromatic discrimination thresholds depended on the contrast of the surround (Silvasti, Valkonen, and Nokelainen 2021), which resembles chromatic contrast induction (Brown and MacLeod 1997) and is simulated by the SBL model. Shadow-illusion effects have also been demonstrated in fish (Simpson, Marshall, and Cheney 2016). Aside from predicting colour appearance the SBL model highlights comparatively unexplored trade-offs in visual systems, with contrast sensitivity potentially linked to dynamic range and to other factors such as low-light vision and temporal acuity. For example, birds have poor luminance contrast sensitivity, but high temporal acuity consistent with a low neural bandwidth in the SBL model (Potier, Mitkus, and Kelber 2018; Ghim and Hodos 2006; Boström et al. 2016).

## Supporting information

Supplementary model code

## Acknowledgements

We thank Jenny Bosten, Nick Scott-Samuel and Roland Baddeley for their constructive feedback.

## Contributions

JT conceived and developed the initial model, and performed the coding and testing; DO contributed to further model development and testing. Both authors wrote the manuscript.

## Supplementary Material

### Description of the Spatiochromatic Bandwidth Limited model

#### Model input Requirements

- A linear cone-catch image of known angular width. For example, cone-catch images created by the micaToolbox (Troscianko & Stevens 2015), or sRGB images converted to linear CIE XYZ channels. Our implementation accepts either sRGB images or cone catch images, and uses 32-bit images and processing throughout; 8-bits per channel is an insufficient dynamic range for coding linear natural scenes. The image should be scaled so that its resolution matches or exceeds the highest spatial frequency being modelled. For example, the DoG kernel we use has its peak wavelength sensitivity at 5.7 pixels, and the highest SF we model is 16 cpd, so the image should be scaled so that each degree of angular width has 16 × 5.7 pixels, i.e. 91.2 pixels per degree.
- Contrast sensitivity functions (CSFs) for the luminance and chromatic opponent channels (red-green and blue-yellow). Our code uses values from Kim et al. (2013). These values should be scaled so that contrasts are Michelson Contrast values - e.g. (red-green)/(red+green). Note that sensitivity is the inverse of the threshold contrast (i.e. *higher* sensitivity = *lower* threshold contrasts, Figure 2a).
- Bandwidth values (ε) for luminance and each chromatic opponent channel (i.e. three values for human vision). These can be estimated from behavioural data (e.g. crispening effect, Figure 3a), or from neurophysiological data (Figure 3b). Suitable data are currently lacking for chromatic channel bandwidth, but we assume the red-green channel bandwidth equals that for the luminance channel, and the blue-yellow channel has about 30% of this bandwidth, in order to achieve efficient coding in natural scenes.
- Gain functions specify how each spatial frequency should be scaled following the clipping process. These are calculated by processing a library of images of natural scenes through the model with all gain values set to 1 (i.e. no gain), and measuring the resulting standard deviation of each channel. Normalising to these values gives output contrasts with standard deviations of 1 at each spatial frequency.

The values we used are supplied in the supplementary code.

#### Image Pre-Processing

The cone catch image is converted to three channels: luminance, red-green opponency and blue-yellow opponency. The luminance channel is the average of all cone catch values from each receptor class, weighted by their cone ratios:

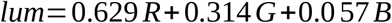

Where R, G, and B are the longwave, mediumwave and shortwave cone catch pixel values respectively. Cone ratios here are from Hofer *et al*. (2005).

The chromatic signals are calculated as Michelson contrasts:

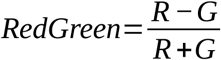

#### Spatial Filtering

Each channel is convolved with either a Difference-of-Gaussian kernel or Gabor kernel. DoG kernels are orientation-insensitive, and are used for luminance and chromatic channels. Gabor kernels are orientation sensitive and are optionally used instead of DoG for the luminance channel. Our implementation uses conventional kernel functions (see code for exact parameters, examples shown in Figure 1b); for the DoG the surround has a sigma value 1.6 times larger than the centre, and for the Gabor filter we use 4 orientations (sigma = 2, gamma = 1, frequency = 3). Our spatial filtering differs from that used previously in that we use Michelson Contrasts. Conventionally, the spatial filtering procedure uses logged input images and then applies a convolution. The result is mathematically identical to dividing the centre response by the surround. While this is computationally efficient, the resulting contrasts are non-linear and unbounded (e.g. values can easily go implausibly high), and cannot be reliably matched to the behaviour described in CSFs. The chromatic channels have already had the Michelson contrast function applied, so the convolution is equivalent to simulating Michelson contrasts based on red-centre versus green-surround, or yellow-centre versus blue-surround giving contrast values, φ. However, the Michelson contrast stage must be applied to the luminance channel following spatial filtering in order to compare centre and surround (or positive and negative regions in the Gabor kernel), i.e.:

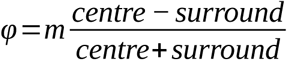

Our implementation achieves this by calculating both signed and unsigned (absolute) convolutions for the numerator and denominator respectively. *m* is a parameter that scales the kernel’s (arbitrary) amplitude to create contrasts that match the same scale as the contrast sensitivity functions. CSFs are generally calculated using sinewave gratings, so to calculate *m* we first create an image with a sinewave spatial frequency that matches the kernel’s peak sensitivity (5.7 pixels in our case). The sinewave amplitude is set to a known Michelson contrast of e.g. 0.1, and then is convolved with the kernel. *m* is then the maximum contrast from the convolved image divided by the Michelson contrast of the input sinewave (i.e. 0.1). This simply scales the contrasts, φ, so that they are directly comparable to the conditions used to measure CSFs.

#### Clipping

The activation threshold, α, is the inverse of contrast sensitivity, specified by the CSF at each spatial frequency, ϖ (see Figure 2a for example CSFs):

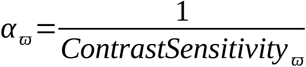

Any contrasts below the saturation threshold are set to zero, while all other contrasts have the saturation threshold subtracted (Kulikowski 1976):

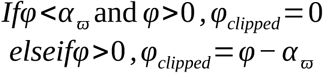

The sign is preserved for negative contrasts (i.e. the model assumes both centre-on and centre-off behaviour, described by positive or negative convolved pixel values respectively). However, in natural scenes the luminance DoG convolution results in negative contrasts that are twice as large as the positive ones (this does not apply to chromatic DoG or Gabor convolutions, where positive contrasts match negative contrasts). Following the principles of efficient coding we therefore assume that any centre-off channels are tuned to the same dynamic range, and multiply α by 2 i.e.:

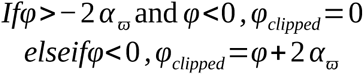

Bandwidth, ε, is assumed to be uniform across all spatial frequencies, and this is used to calculate the saturation threshold, β, at each spatial frequency:

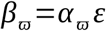

The bandwidth can either be estimated by fitting the model to behavioural data (Figure 3a), or based on the dynamic range of single neurones (Figure 3b). Contrasts greater than the saturation threshold are set to equal the saturation threshold, creating a hard upper threshold. As above, negative contrasts are doubled for luminance DoG models (but not chromatic or Gabor models):

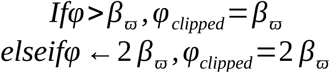

This clipping process defines the dynamic range of the model at each spatial frequency. The result is that spatial frequencies with high contrast sensitivity also saturate much faster with increasing contrast, resulting in small dynamic range. Meanwhile spatial frequencies with low contrast sensitivity have a much larger dynamic range (Figure 2b). However, the overlap in sensitivity between adjacent spatial frequencies means that almost all contrasts are within the dynamic range of one or more spatial frequencies (the orange areas in Figure 2f), implying low bandwidths can be combined with high contrast sensitivity for efficient coding as long as there is a large range in dynamic ranges, and sufficient overlap in adjacent spatial frequencies. This explains why humans can perceive contrasts in natural scenes (or on high-definition televisions) over a dynamic range greater than 10,000:1, while our dynamic range for sinewaves is around 200:1.

**Supplementary Figure 1.**
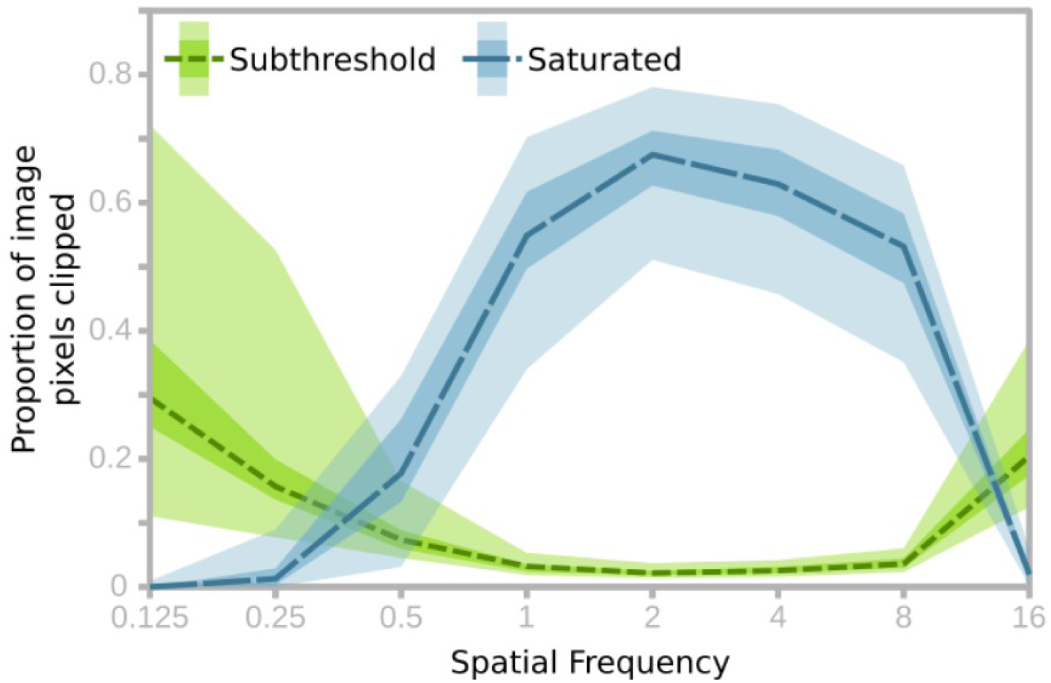
Plot showing the proportion of pixels in each channel that are either saturated or sub-threshold in typical natural scenes. The results are based on 34 images of natural scenes, dashed lines show the median value, shaded areas show the interquartile range and full range of the data. High contrast sensitivity at intermediate spatial frequencies causes substantially more saturation, while the lower sensitivity channels show substantial proportion of subthreshold contrasts.

#### Gain

Following clipping, contrasts are multiplied so that each spatial frequency results in equal contribution to the contrasts in the pooled image. i.e. in natural scene statistics each spatial frequency should contain equal contrast/information (Field, 1987), however the clipping process substantially reduces the average amplitude of contrasts at intermediate spatial frequencies. The gain step equalises the average contrast amplitudes at each spatial frequency, i.e.:

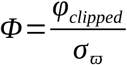

Where σ_ϖ_ is the standard deviation of all φ_clipped_ values in an image of a natural scene filtered at spatial frequency ϖ, resulting in gain-corrected contrasts, Φ.

#### Post-clipping smoothing

The hard upper and lower clipping thresholds (α and β) produce undesirable artefacts in the pooled image. We remove these by applying a Gaussian blur to each channel prior to pooling, with sigma values well below the filter’s spatial frequency (e.g. sigma value below 1 pixel radius, where the kernel’s peak wavelength sensitivity is 5.7 pixels). This step removes the artefacts, and the smoothing effect is responsible for the curvature near the saturation threshold shown in Figure 3b, matching the behaviour of primate ganglia (Derrington and Lennie, 1984). This stage mirrors the correlated firing of neighbouring retinal ganglion cells cells where on-centre cells excite neighbouring on-centre cells, and likewise for off-centre cells, while on-centre and off-centre cells inhibit one-another (Nelson 1995).

#### Pooling

Pooling simply sums the contrast at each pixel location across each spatial frequency:

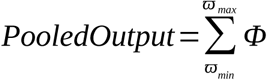

This results in recombined luminance, red-green and blue-yellow chromatic channels. This output is designed to match subjective colour appearance, and it is therefore not straightforward to present these images on an sRGB display without confounding the very effects it seeks to predict. Nevertheless, we can convert back to a space that roughly approximates the cone-catch input:

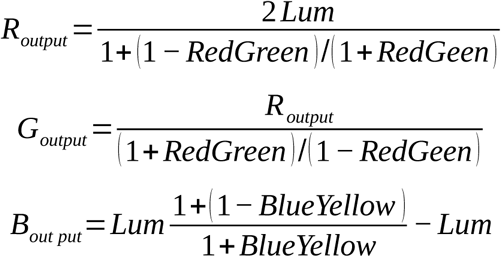

## Appendix to

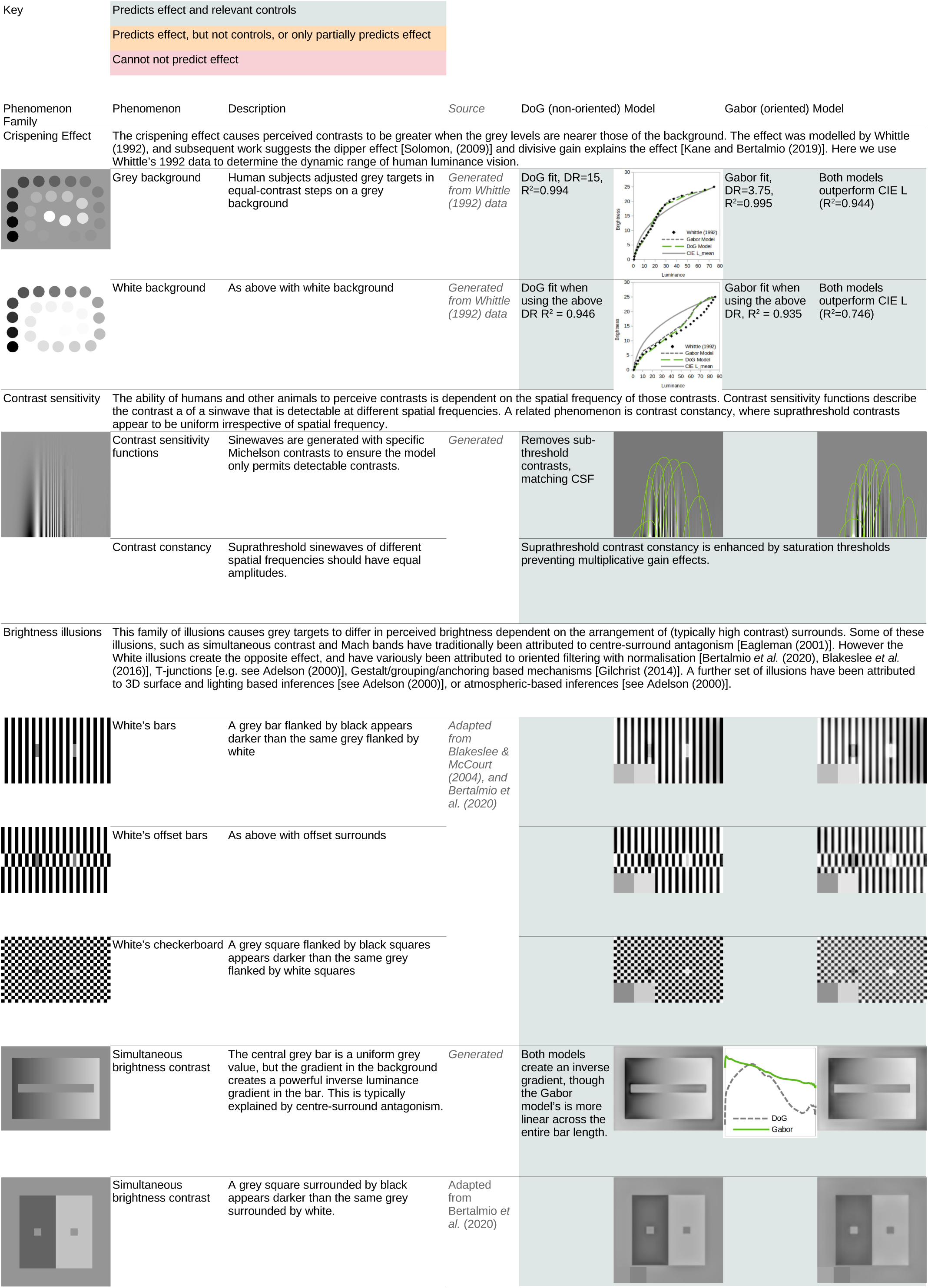

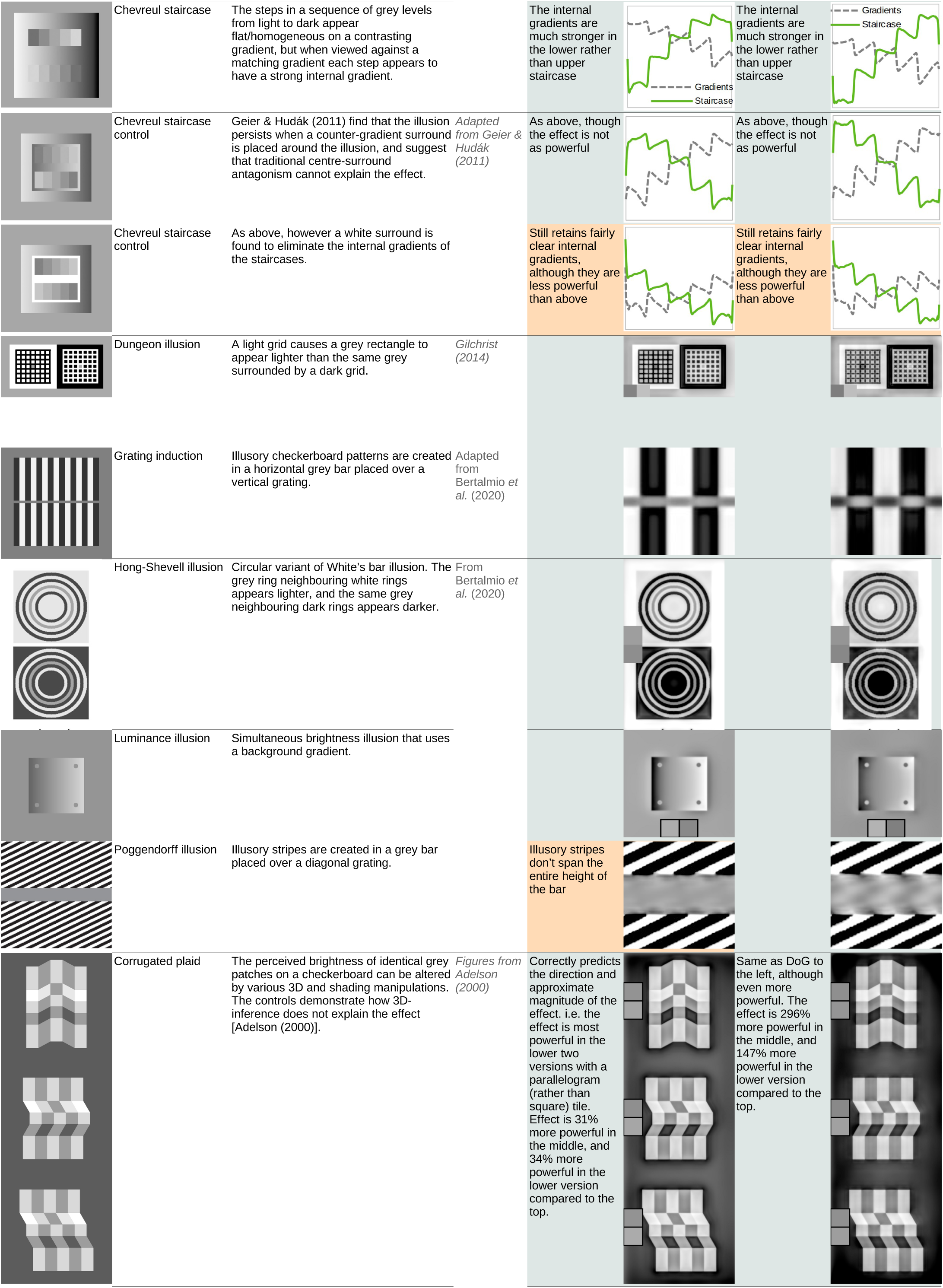

|  |  |  |  |  |  |  |
| --- | --- | --- | --- | --- | --- | --- |
|  | Chevreul staircase | The steps in a sequence of grey levels from light to dark appear flat/homogeneous on a contrasting gradient, but when viewed against a matching gradient each step appears to have a strong internal gradient. |  | The internal gradients are much stronger in the lower rather than upper staircase |  | The internal gradients are much stronger in the lower rather than upper staircase |
|  | Chevreul staircase control | Geier & Hudák (2011) find that the illusion persists when a counter-gradient surround is placed around the illusion, and suggest that traditional centre-surround antagonism cannot explain the effect. | *Adapted from Geier & Hudák (2011)* | As above, though the effect is not as powerful |  | As above, though the effect is not as powerful |
|  | Chevreul staircase control | As above, however a white surround is found to eliminate the internal gradients of the staircases. |  | Still retains fairly clear internal gradients, although they are less powerful than above |  | Still retains fairly clear internal gradients, although they are less powerful than above |
|  | Dungeon illusion | A light grid causes a grey rectangle to appear lighter than the same grey surrounded by a dark grid. | *Gilchrist (2014)* |  |  |  |
|  | Grating induction | Illusory checkerboard patterns are created in a horizontal grey bar placed over a vertical grating. | *Adapted from Bertalmio et al. (2020)* |  |  |  |
|  | Hong-Shevell illusion | Circular variant of White's bar illusion. The grey ring neighbouring white rings appears lighter, and the same grey neighbouring dark rings appears darker. | *From Bertalmio et al. (2020)* |  |  |  |
|  | Luminance illusion | Simultaneous brightness illusion that uses a background gradient. |  |  |  |  |
|  | Poggendorff illusion | Illusory stripes are created in a grey bar placed over a diagonal grating. |  | Illusory stripes don't span the entire height of the bar |  |  |
|  | Corrugated plaid | The perceived brightness of identical grey patches on a checkerboard can be altered by various 3D and shading manipulations. The controls demonstrate how 3D-inference does not explain the effect [Adelson (2000)]. | *Figures from Adelson (2000)* | Correctly predicts the direction and approximate magnitude of the effect. i.e. the effect is most powerful in the lower two versions with a parallelogram (rather than square) tile. Effect is 31% more powerful in the middle, and 34% more powerful in the lower version compared to the top. |  | Same as DoG to the left, although even more powerful. The effect is 296% more powerful in the middle, and 147% more powerful in the lower version compared to the top. |

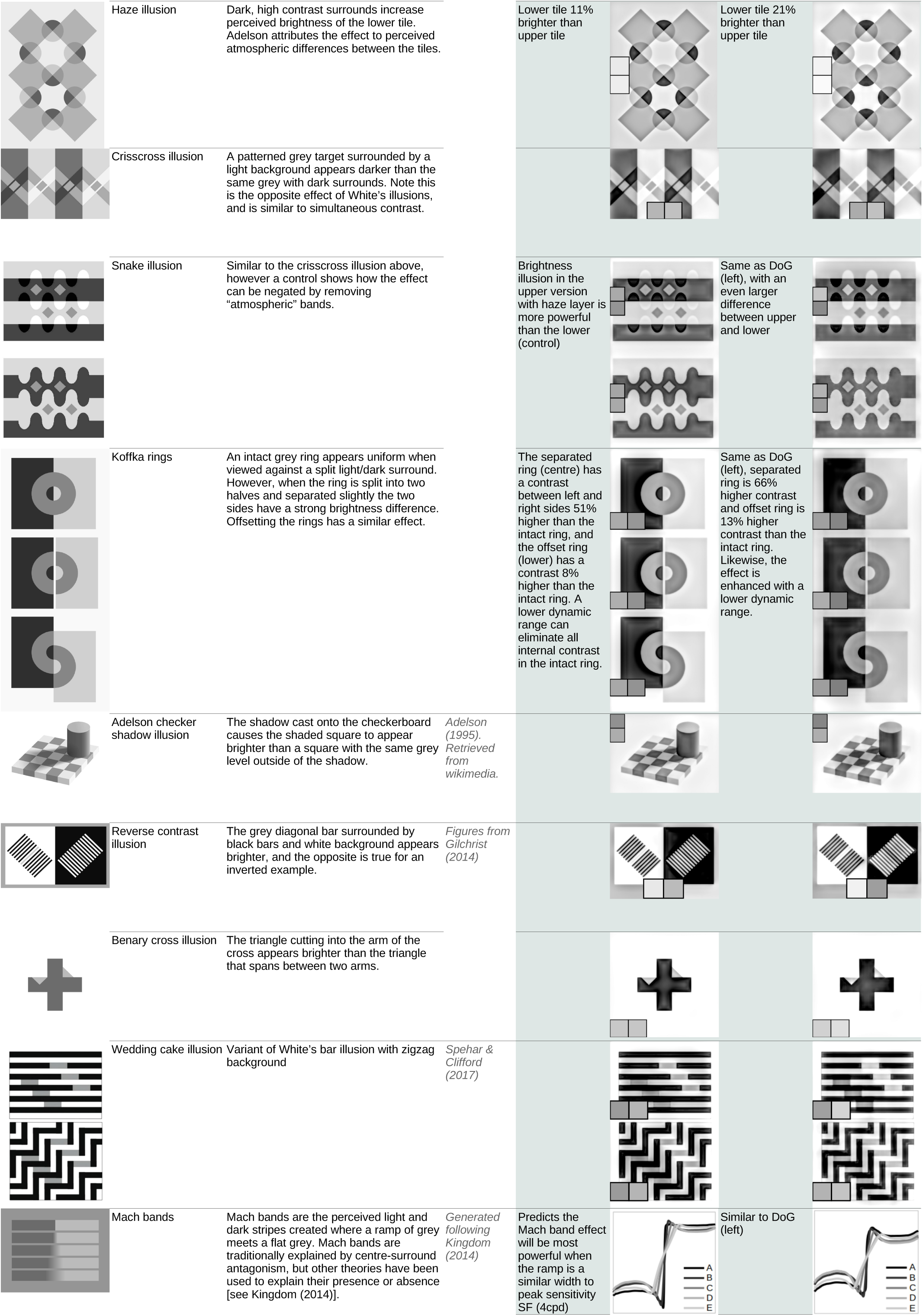

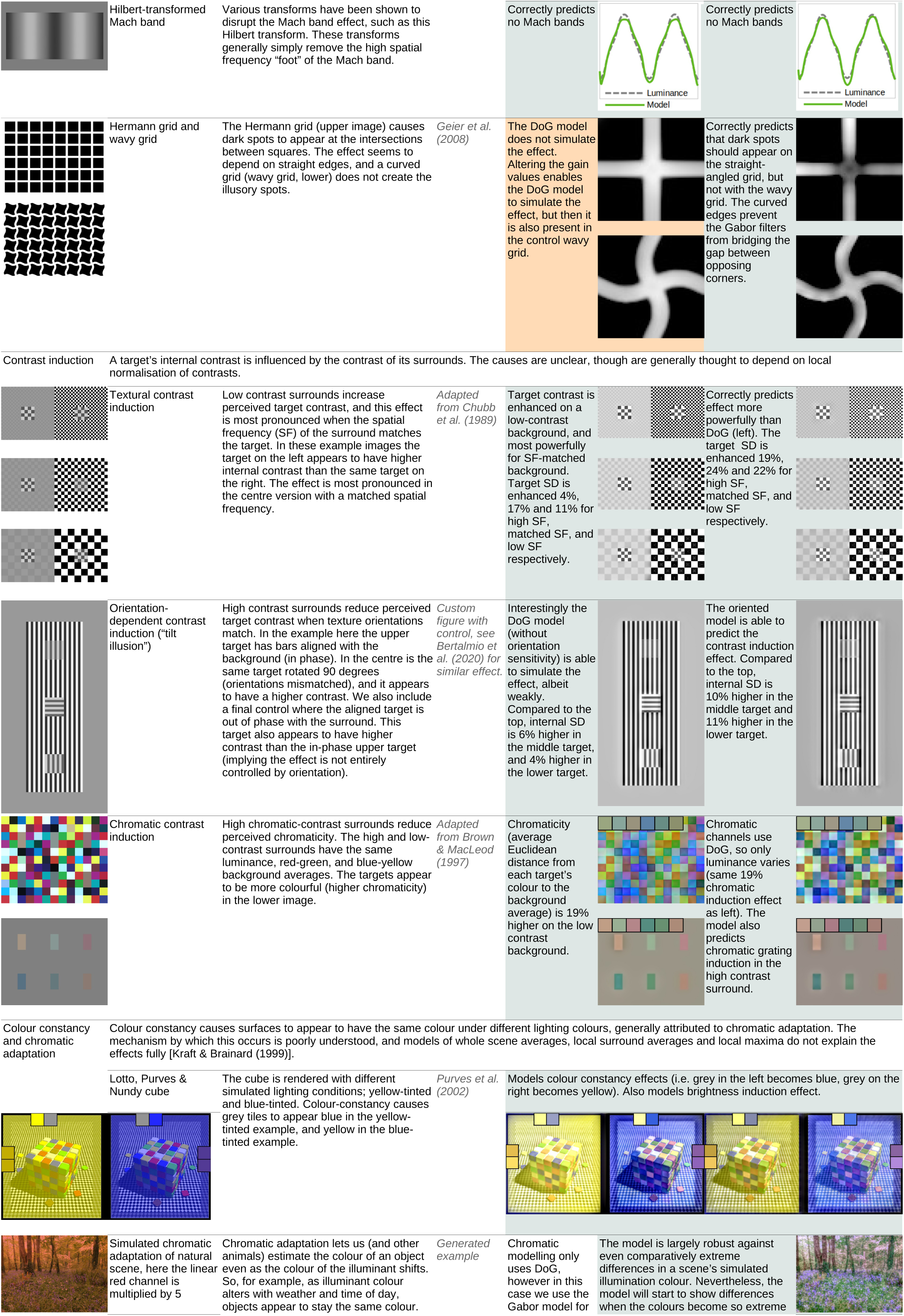

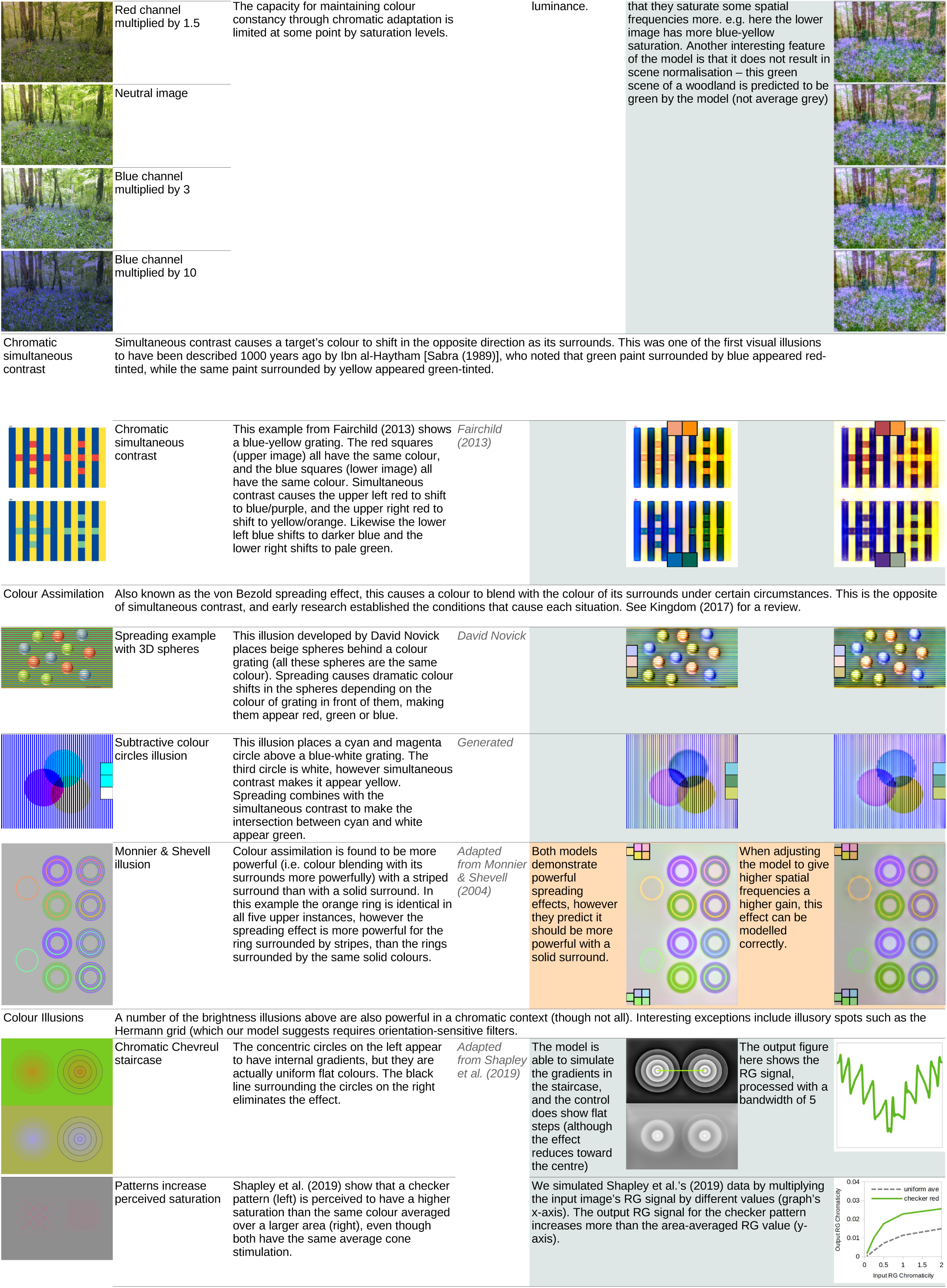

|  |  |  |  |  |
| --- | --- | --- | --- | --- |
|  | Red channel multiplied by 1.5 | The capacity for maintaining colour constancy through chromatic adaptation is limited at some point by saturation levels. | luminance. | that they saturate some spatial frequencies more, e.g. here the lower image has more blue-yellow saturation. Another interesting feature of the model is that it does not result in scene normalisation – this green scene of a woodland is predicted to be green by the model (not average grey) |
|  | Neutral image |  |  |  |
|  | Blue channel multiplied by 3 |  |  |  |
|  | Blue channel multiplied by 10 |  |  |  |

|  |  |  |  |
| --- | --- | --- | --- |
|  | Chromatic simultaneous contrast | This example from Fairchild (2013) shows a blue-yellow grating. The red squares (upper image) all have the same colour, and the blue squares (lower image) all have the same colour. Simultaneous contrast causes the upper left red to shift to blue/purple, and the upper right red to shift to yellow/orange. Likewise the lower left blue shifts to darker blue and the lower right shifts to pale green. | *Fairchild (2013)* |

|  |  |  |  |
| --- | --- | --- | --- |
|  | Spreading example with 3D spheres | This illusion developed by David Novick places beige spheres behind a colour grating (all these spheres are the same colour). Spreading causes dramatic colour shifts in the spheres depending on the colour of grating in front of them, making them appear red, green or blue. | *David Novick* |
|  | Subtractive colour circles illusion | This illusion places a cyan and magenta circle above a blue-white grating. The third circle is white, however simultaneous contrast makes it appear yellow. Spreading combines with the simultaneous contrast to make the intersection between cyan and white appear green. | *Generated* |

|  |  |  |  |  |  |  |
| --- | --- | --- | --- | --- | --- | --- |
|  | Monnier & Shevell illusion | Colour assimilation is found to be more powerful (i.e. colour blending with its surrounds more powerfully) with a striped surround than with a solid surround. In this example the orange ring is identical in all five upper instances, however the spreading effect is more powerful for the ring surrounded by stripes, than the rings surrounded by the same solid colours. | *Adapted from Monnier & Shevell (2004)* | Both models demonstrate powerful spreading effects, however they predict it should be more powerful with a solid surround. |  | When adjusting the model to give higher spatial frequencies a higher gain, this effect can be modelled correctly. |

|  |  |  |  |  |  |  |  |
| --- | --- | --- | --- | --- | --- | --- | --- |
|  | Chromatic Chevreul staircase | The concentric circles on the left appear to have internal gradients, but they are actually uniform flat colours. The black line surrounding the circles on the right eliminates the effect. | *Adapted from Shapley et al. (2019)* | The model is able to simulate the gradients in the staircase, and the control does show flat steps (although the effect reduces toward the centre) |  |  | The output figure here shows the RG signal, processed with a bandwidth of 5 |
|  | Patterns increase perceived saturation | Shapley et al. (2019) show that a checker pattern (left) is perceived to have a higher saturation than the same colour averaged over a larger area (right), even though both have the same average cone stimulation. |  | We simulated Shapley et al.'s (2019) data by multiplying the input image's RG signal by different values (graph's x-axis). The output RG signal for the checker pattern increases more than the area-averaged RG value (y-axis). |  |  |  |

