## Supplementary model code for "A model of colour appearance based on efficient coding of natural images": DoG_ScC_Model_Michelson.java

```
import ij.*;
import ij.process.*;
import ij.gui.*;
import java.awt.*;
import ij.plugin.*;
import ij.util.*;

import ij.measure.ResultsTable;

/*

	Title: DoG ROI Bandpass
	Author: Jolyon Troscianko
	Date: 17/9/2021

.................................................................................................................

Description:
'''''''''''''''''''''

This script applies a bank of DoG filters to an ROI. The script has been designed to
work with ROIs of any (non-rectangular) shape by applying a 2D convolution and
weighting all measurements by the degree to which the kernel is inside the ROI
i.e. it re-normalises the kernel with each convolution.

The script efficiently applies the DoG filter to a series of different spatial scales
by effectively scaling the size of the image (with no interpolation). This makes
it incredibly fast at measuring the larger spatial frequencies that would otherwise
be incredibly slow to process (e.g. by using a larger sigma value).

The script then applies a bicubic interpolation (Catmull-Rom) to the output. This
interpretation assumes that pixels outside the ROI have a value of zero, which
makes sense given the convolution is normalised to have average output of zero.
This is different from a standard image, where out-of-bounds pixels cannot be assumed
to have a particular value.

The result is a very fast convolution, with increasing processing efficiency at larger
spatial scales, combined with smooth output. Any systematic errors introduced by
the scaling and interpolation will be applied equally at all spatial scales.


Input Requirements:
'''''''''''''''''''''
Requires a 32-bit image. If an ROI is selected it will use it, otherwise it will
process the whole image.

The size of the selection must be larger than 2^nOctaves otherwise
the entire image will be a single pixel at larger scales. If the SD ("energy")
is zero at a given scale then avoid using that scale or greater, and probably
avoid using the first smaller scale.

*/
public class DoG_ScC_Model_Michelson implements PlugIn {
public void run(String arg) {


double negM = 2.0; // in a natural scene, unclipped, the negative median is almost exacly 2 times larger than the positive (for all but lowes SFs where sampling error comes in)

double DoGMmichelson = 2.75; // 2.75
double DoGM = 0.06; // approx 0.04 cuts off contrast


/*
calculating DoG multiplier: 

This metric is purely designed to match the way contrast is measured in CSF measurements.
Switch off clipping and weighting before doing this testing (all gains =1)!

----Lum testing with michelson contrast----------
Input image Michelson contrast =0.099, model set ot no clipping, equal gains,
at the SF associated with 16 cpd (around 5 pixels wavelength)
the max distance from zero is around 0.008. 0.099/0.008 = 12.37


---Chrom testing (no michelson contrast for the kernel----------
Input image Michelson contast = 0.01, otehrwise as above. Max dist from zero
at around 5px wavelength = 0.05. 0.01/0.05 = 0.2

The highest SF (16cpd) has a peak SF of 8 pixels wavelength (much lower frequ than Gabor)
It does have a slightly higher amplitude response to longer wavelengths, but this freq will give
the highest energy response

So a 19 degree wide image would be 2432 pixels wide


*/


//-----------------CS Values from Kim et al 2013----------------------
//double[]  lumSF = {0.125,0.25,0.5,1,2,4,8,16};
//double[] lumCS = {21.77,39.98,86.14,197.83,266.64,229.67,181.65,37.91};
// order matching k
//double[] lumCS = {37.91, 181.65, 229.67, 266.64, 197.83, 86.14, 39.98, 21.77};


// First crispening itteration fitting weights based on DR216 PS1.75 using linear image of woodland 1730px wide
//double[] lumWeights = {0.7722885594230237,0.3221101597276327,0.2665152034754664,0.24201084046593935,0.2907966514020694,0.41970275026166337,0.41804969782160006,0.7650995725917734};

// as above, but measured using a central selection area away from edges (edge effect etc..)
//double[] lumWeights = {0.663619854, 0.306713962, 0.261508905, 0.222250459, 0.261638294, 0.379001169, 0.348794227, 0.472771658};


	ImagePlus imp = IJ.getImage();

	// ---------------------- Calculate Gabor Filter Stuff-------------------------------

	double sigma1 = 1.0;
	double sigma2 = 1.6;
	//int nOctaves = 8;
	String imTitle = imp.getTitle();
	double dr = 15;
	double postSigma = 0.75;
	double poolingWeight = 1.0;


	GenericDialog gd = new GenericDialog("DoG Convolution");
		gd.addMessage("Wavelet Filter Settings");
		gd.addNumericField("Sigma1", sigma1, 2);
		gd.addNumericField("Sigma2", sigma2, 2);
		gd.addCheckbox("Michelson", true);
		gd.addCheckbox("Clipping", true);
		gd.addNumericField("Dynamic range", dr, 2);
		gd.addNumericField("Post threshold sigma", postSigma, 2);
		//gd.addNumericField("Number_of_octaves", nOctaves, 0);
		gd.addStringField("Spatial_frequency", "0.125,0.25,0.5,1,2,4,8,16", 50);
		//gd.addStringField("CSF", "21.77,39.98,86.14,197.83,266.64,229.67,181.65,37.91", 50); // values from kim et al 2016 using relative modulation
		// LUM
		gd.addStringField("CSF", "10.885,19.99,43.07,98.915,133.32,114.835,90.825,18.955", 50); // values from kim et al 2016 divided by two, for use with michelson contrast
	
		// RG:
		//gd.addStringField("CSF", "307.3,344.8,307.3,261.5,154.0,97.2,0,0", 50); // Kim et al 2013 values converted from their "C" number to michelson contrast. See spreadsheet for conversion

		//gd.addStringField("Gain", "0.671,0.49,0.540,0.383,0.334,0.386,0.456,1.05", 50);
		//gd.addStringField("Gain", "0.381,0.298,0.253,0.147,0.113,0.128,0.161,0.494", 50); // DR16.5 PS0.8 2nd itteration weights
		//gd.addStringField("Gain", "0.175,0.186,0.186,0.125,0.099,0.113,0.140,0.369", 50); // DR15 PS0.8 DoGMMich 2.75 3rd itteration weights
		gd.addStringField("Gain", "0.201,0.221,0.179,0.097,0.074,0.087,0.109,0.309", 50); // DDR9.5 PS0.6 DoGMMich 2.75 Natural scene database
		//gd.addStringField("Gain", "0.205,0.244,0.261,0.181,0.147,0.170,0.206,0.367", 50); // DDR20 PS0.6 DoGMMich 2.75 Natural scene database
		//gd.addStringField("Gain", "0.186,0.218,0.211,0.130,0.103,0.120,0.149,0.332", 50); // DR15 PS0.75 DoGMMich 2.75 Natural scene database

		//gd.addStringField("Gain", "1,1,1,1,1,1,1,1", 50);
		gd.addNumericField("Pooling weight", poolingWeight, 2);
		gd.addCheckbox("Interpolation", true);
		gd.addMessage("Output Images");
		gd.addCheckbox("DoG Kernel", false);
		gd.addStringField("Label", imTitle, 20);

	gd.showDialog();
	if (gd.wasCanceled())
		return;


	sigma1 = gd.getNextNumber();
	sigma2 = gd.getNextNumber();
	boolean michelson = gd.getNextBoolean();

	if(michelson == false)
		negM = 1.0;

	boolean clipping = gd.getNextBoolean();
	dr = gd.getNextNumber();
	postSigma = gd.getNextNumber();
	//nOctaves = (int) Math.round(gd.getNextNumber());

// Import input data
	String[] lumSFstring = Tools.split(gd.getNextString(), ",");
	//----------convert string to numeric array-----------
	double[] lumSF = new double[lumSFstring.length];
	double peakSF = 0.0;
	for(int i=0; i<lumSFstring.length; i++){
		lumSF[i] = Double.valueOf(lumSFstring[i]);
		if(lumSF[i] > peakSF)
			peakSF = lumSF[i]; // save peak for use in calculating the scale
	}
	

	String[] lumCSstring = Tools.split(gd.getNextString(), ",");
	//----------convert string to numeric array-----------
	double[] lumCS = new double[lumCSstring.length];
	for(int i=0; i<lumCSstring.length; i++){
		lumCS[i] = Double.valueOf(lumCSstring[i]);
		//IJ.log("SF: " + lumSF[i] + " CS: " + lumCS[i]);
	}

	String[] lumWeightsstring = Tools.split(gd.getNextString(), ",");
	//----------convert string to numeric array-----------
	double[] lumWeights = new double[lumWeightsstring.length];
	for(int i=0; i<lumWeightsstring.length; i++)
		lumWeights[i] = Double.valueOf(lumWeightsstring[i]);
		
	if(lumWeights.length != lumCS.length || lumWeights.length != lumSF.length){
		IJ.log("Error - input value lengths don't match, quitting");
		return;
	}

	int nOctaves = lumCS.length;

	poolingWeight = gd.getNextNumber();
	boolean interpolate = gd.getNextBoolean();
	boolean outputKernel = gd.getNextBoolean();

	imTitle = gd.getNextString();


//---normalise gain/weights array to mean=1------

double meanLumWeight = 0.0;
for(int i=0; i<lumWeights.length; i++)
	meanLumWeight += lumWeights[i];

meanLumWeight = meanLumWeight/lumWeights.length;

for(int i=0; i<lumWeights.length; i++)
	lumWeights[i] = lumWeights[i]/meanLumWeight;

for(int i=0; i<lumWeights.length; i++)
	lumWeights[i] = lumWeights[i]*poolingWeight + (1-poolingWeight);


// sigma 2 should be larger than sigma1
if(sigma1 > sigma2){
	double td = sigma2;
	sigma2 = sigma1;
	sigma1 = td;
}

//----------------calculate size of DoG kernel---------------

double gVal = (1/Math.pow(2*Math.PI*Math.pow(sigma2,2),0.5)) * Math.exp(-1*(Math.pow(0,2)/(2*Math.pow(sigma2,2) ) ));
double minVal = 0.001; // this is the cutoff as a proportion of the peak at which it is assumed the Guassian curve is small enough to have no influence
minVal = minVal / ( (1/Math.pow(2*Math.PI*Math.pow(sigma2,2),0.5)) * Math.exp(-1*(Math.pow(0,2)/(2*Math.pow(sigma2,2) ) ))); // minVal as a proportaion of start val
int cutoff = 1;

while(gVal > minVal){
	gVal = (1/Math.pow(2*Math.PI*Math.pow(sigma2,2),0.5)) * Math.exp(-1*(Math.pow(cutoff,2)/(2*Math.pow(sigma2,2) ) ));
	cutoff++;
}


// Gaussian Kernel
int mapW = 2*cutoff +1;
double[] kernelArray = new double[mapW*mapW]; 
double[] absKernelArray = new double[mapW*mapW];
double rad = 0.0;

// create normalised kernel where mean = 0
double cSum = 0.0; // centre sum
double sSum = 0.0; // surround sum
double csSD = 0.0; // sum of squares  for SD

//---- Calculate the area under the curve for centre and surround Gaussian sigma value for normalisation

for(int y=0; y<mapW; y++)
for(int x=0; x<mapW; x++){
	rad = Math.pow( Math.pow(x-cutoff,2) + Math.pow(y-cutoff,2) ,0.5);
	sSum += (1/Math.pow(2*Math.PI*Math.pow(sigma2,2),0.5)) * Math.exp(-1*(Math.pow(rad,2)/(2*Math.pow(sigma2,2) ) ));
	cSum += (1/Math.pow(2*Math.PI*Math.pow(sigma1,2),0.5)) * Math.exp(-1*(Math.pow(rad,2)/(2*Math.pow(sigma1,2) ) ));
}


for(int y=0; y<mapW; y++)
for(int x=0; x<mapW; x++){
	rad = Math.pow( Math.pow(x-cutoff,2) + Math.pow(y-cutoff,2) ,0.5);
	kernelArray[(y*mapW)+x] = -((1/Math.pow(2*Math.PI*Math.pow(sigma2,2),0.5)) * Math.exp(-1*(Math.pow(rad,2)/(2*Math.pow(sigma2,2) ) )))/sSum;
	kernelArray[(y*mapW)+x] += ((1/Math.pow(2*Math.PI*Math.pow(sigma1,2),0.5)) * Math.exp(-1*(Math.pow(rad,2)/(2*Math.pow(sigma1,2) ) )))/cSum;
	csSD += Math.pow(kernelArray[(y*mapW)+x] , 2);
}

// ---- normalise DoG Kernel so that SD=1---------

csSD = Math.pow(csSD/(mapW*mapW), 0.5);

for(int y=0; y<mapW; y++)
for(int x=0; x<mapW; x++){
	kernelArray[(y*mapW)+x] = kernelArray[(y*mapW)+x] / csSD;
	absKernelArray[(y*mapW)+x] = Math.abs(kernelArray[(y*mapW)+x]);
}


	if(outputKernel == true){
		ImageStack kernelStack = new ImageStack(mapW, mapW);
		float[] kernelOutputArray = new float[mapW*mapW];
		for(int i=0; i<mapW*mapW; i++)
			kernelOutputArray[i] = (float) kernelArray[i];
		kernelStack.addSlice("s" + sigma1 + "_" +sigma2, kernelOutputArray);
		new ImagePlus("DoG Kernel", kernelStack).show();
	}


//----------------calculate size of Gaussian (post-clipping) kernel---------------

double pgVal = (1/Math.pow(2*Math.PI*Math.pow(postSigma,2),0.5)) * Math.exp(-1*(Math.pow(0,2)/(2*Math.pow(postSigma,2) ) ));
double pminVal = 0.001; // this is the cutoff as a proportion of the peak at which it is assumed the Guassian curve is small enough to have no influence
pminVal = pminVal / ( (1/Math.pow(2*Math.PI*Math.pow(postSigma,2),0.5)) * Math.exp(-1*(Math.pow(0,2)/(2*Math.pow(postSigma,2) ) ))); // minVal as a proportaion of start val
int pcutoff = 1;

while(pgVal > pminVal){
	pgVal = (1/Math.pow(2*Math.PI*Math.pow(postSigma,2),0.5)) * Math.exp(-1*(Math.pow(pcutoff,2)/(2*Math.pow(postSigma,2) ) ));
	pcutoff++;
}

// Gaussian Kernel
int pmapW = 2*pcutoff +1;
double[] pgausMap = new double[pmapW*pmapW]; 
double prad = 0.0;

for(int y=0; y<pmapW; y++)
for(int x=0; x<pmapW; x++){
	prad = Math.pow( Math.pow(x-pcutoff,2) + Math.pow(y-pcutoff,2) ,0.5);
	pgausMap[(y*pmapW)+x] = (1/Math.pow(2*Math.PI*Math.pow(postSigma,2),0.5)) * Math.exp(-1*(Math.pow(prad,2)/(2*Math.pow(postSigma,2) ) ));
}


// --------------------- Get ROI Mask and Outline---------------------------------


	Roi roi = imp.getRoi();
	if (roi!=null && !roi.isArea()) roi = null;
	ImageProcessor ip = imp.getProcessor();
	int w = ip.getWidth();
	int h = ip.getHeight();
	int roiW = 0;
	int roiH = 0;
	int roiX = 0;
	int roiY = 0;

	//ImageProcessor mask = new ByteProcessor(roiW, roiH);

	if (roi==null) // in case there is no ROI selected use the entire image
		roi = new Roi(0, 0, w, h);
	
	Rectangle r = roi.getBounds();
	ImageProcessor mask = roi.getMask();

	//int roiType = roi.getType();
	//if (roiType==Roi.RECTANGLE){
	if(mask==null){
		//IJ.log("rectangle");
		mask = new ByteProcessor(r.width, r.height);
		mask.invert();
	}

	roiW = r.width;
	roiH = r.height;
	roiX = r.x;
	roiY = r.y;


ImageStack outStack = new ImageStack(roiW, roiH);
double kernelSum = 0.0; // the kernel needs re-normalising with each pass
int kernelCount = 0;
double absKernelSum = 0.0;
double convSum = 0.0;
double absConvSum = 0.0;
double rawSum = 0.0;
double tpx = 0.0;


float[] pixels = new float[roiW*roiH];

for (int y=0; y<roiH; y++)
for (int x=0; x<roiW; x++)
if(mask.getPixel(x,y) != 0){
	pixels[x+ y*roiW] = ip.getPixelValue(x+roiX, y+roiY);
}


//ResultsTable rt =  ResultsTable.getResultsTable();
//rt.incrementCounter();
//rt.addValue("Title", imTitle);


//---------------Bicubic interpolation coefficients----------------
double[][] M = {             // Catmull-Rom basis matrix
	{-0.5,  1.5, -1.5, 0.5},
	{ 1  , -2.5,  2  ,-0.5},
	{-0.5,  0  ,  0.5, 0  },
	{ 0  ,  1  ,  0  , 0  }
};

double[][] C = new double[4][4];
double[][] T = new double[4][4];
double[][] G = new double[4][4];

double ix = 0.0;
double iy = 0.0;


//-------------------------------- downscale pixels-------------------------------------
// sampling at an octave scale

float[] pooledPxs = new float[roiW*roiH];

double aTpos = 0.0;
double aTneg = 0.0;
double sTpos = 0.0;
double sTneg = 0.0;

int scaleRad = 1;
String ts = "";
String tss = "";
double aveEnergy = 0.0;

String satString = "";


for(int k=0; k<nOctaves; k++)
if(lumCS[k] > 0.0){

//scaleRad = (int) Math.pow(2, nOctaves-k-1);

scaleRad = (int) Math.round(peakSF / lumSF[k]);

int scaleW = (int) (Math.floor(roiW/scaleRad) +1);
int scaleH = (int) (Math.floor(roiH/scaleRad) +1);

double scaleSum = 0.0;
int scaleCount = 0;


float[] scalePixels = new float[scaleW*scaleH];
int[] scaleMask = new int[scaleW*scaleH];

for(int y=0; y<scaleH; y++)
for(int x=0; x<scaleW; x++){

	scaleSum = 0.0;
	scaleCount = 0;

	for(int y2 = 0; y2<scaleRad; y2++)
	for(int x2 = 0; x2<scaleRad; x2++)
	if(mask.getPixel(x2+x*scaleRad,y2+y*scaleRad) != 0){
		scaleSum += pixels[(x2+x*scaleRad) + (roiW*(y2+y*scaleRad))];
		scaleCount ++;
	} //x2 y2 fi

	if(scaleCount > 0){
		scalePixels[x+y*scaleW] = (float) (scaleSum/scaleCount);
		scaleMask[x+y*scaleW] = 1;
	}
} //xy


//-----------------Scale convolution-------------


double[] pixels2 = new double[scaleW*scaleH];
int[] counts = new int[scaleW*scaleH];
float[] outPxs = new float[scaleW*scaleH];

int loc = 0;
int pxloc = 0;
int scaleLoc = 0;

for(int y=0; y<scaleH; y++)
for(int x=0; x<scaleW; x++)
if(scaleMask[x + scaleW*y] == 1){

	pxloc = (y*scaleW)+x;

	rawSum = 0.0;
	convSum = 0.0;
	absConvSum = 0.0;
	kernelSum = 0.0;
	absKernelSum = 0.0;
	kernelCount = 0;

	for(int y2=-cutoff; y2<=cutoff; y2++)  
	for(int x2=-cutoff; x2<=cutoff; x2++){
		if(y+y2>=0 && y+y2< scaleH && x+x2 >=0 &&x+x2 < scaleW && scaleMask[x + x2 + scaleW*(y+y2)] == 1){
			loc = ((y2+cutoff)*mapW) + (x2+cutoff);

			tpx = scalePixels[x+x2 + (scaleW*(y+y2))];
			rawSum += tpx;

			kernelSum += kernelArray[loc];
			kernelCount++;
			convSum += tpx * kernelArray[loc];

			if(michelson == true){
				absKernelSum += absKernelArray[loc];
				absConvSum  += tpx * absKernelArray[loc];
			}
		}
	}

	//pixels2[pxloc] = DoGMultiplier * ((convSum-(kernelSum*(rawSum/kernelCount)))  / (absConvSum/2)); // relative modulation, as used by Kim et al 2016 for CSF measures
	if(michelson == true)
		pixels2[pxloc] = DoGMmichelson * ((convSum-(kernelSum*(rawSum/kernelCount)))  / absConvSum); // Michelson contrast
	else
		pixels2[pxloc] = DoGM * (convSum-(kernelSum*(rawSum/kernelCount)));

	IJ.showProgress((float) y/roiH );
	
} // xy fi


//for(int i=0; i<scaleH*scaleW; i++)
//	outPxs[i] = (float) (pixels2[i]/counts[i]);

//scaleOutStack.addSlice("output angle" + j, outPxs);


//-----------------Apply to original scale (allows for easier measurement and weighting----------


float[] scaleOutPxs = new float[roiW*roiH];

for(int i=0; i<roiW*roiH; i++)
	scaleOutPxs[i] = Float.NaN;

double td = 0.0;
double pxSum = 0.0;
double nPx = 0.0;

//for(int i=0; i<scaleW*scaleH; i++)
//	pixels2[i] = pixels2[i]/counts[i];


// -------------------activation and saturation thresholding---------------

if(clipping == true){

int countSat = 0;
int countUnsat = 0;

//aTpos = (1/lumCS[lumCS.length-k-1]); // note that the octave scale order is reversed by k
aTpos = (1/lumCS[k]);
aTneg = aTpos*-negM;
//aTneg = aTpos*-1.0;
sTpos = aTpos*dr; // saturation threshold
sTneg = aTneg*dr;

//IJ.log("SF:" + lumSF[k] + " aT+:" + aTpos + " aT-:" + aTneg + " sT+:" + sTpos + " sT-:" + sTneg);


for(int i=0; i<scaleW*scaleH; i++){

		td = pixels2[i];
		double tvv = 0.0;
		
		if(td > aTpos)
			tvv = td - aTpos;
		else if(td > 0)
			tvv = 0.0;
		else if(td < aTneg)
			tvv = td - aTneg;
		else if(td < 0)
			tvv = 0.0;
		else tvv = 0.0;

		if(tvv > sTpos){
			tvv = sTpos;
			countSat ++;
		} else if(tvv < sTneg){
			tvv = sTneg;
			countSat ++;
		} else {
			countUnsat ++;
		}

		pixels2[i] = tvv/lumWeights[k];
}//i


satString = satString + "," + String.format("%.4f",  ((double) countSat/ (double) (countSat + countUnsat)) );

}// clipping ture


//-------------------------------------------------------------------------------------------
//--------------------------Post-clipping Gaussian smoothing--------------------------------

//IJ.log("pcutoff: " + pcutoff);
//IJ.log("pmapW: " + pmapW);
//IJ.log("pgausMap.length: " + pgausMap.length);

double[] pixels3 = new double[scaleW*scaleH];
double[] counts2 = new double[scaleW*scaleH];

for(int y=0; y<scaleH; y++)
for(int x=0; x<scaleW; x++){
	pxloc = (y*scaleW)+x;
	if(scaleMask[x + scaleW*y] == 1)
	for(int y2=-pcutoff; y2<=pcutoff; y2++)  
	for(int x2=-pcutoff; x2<=pcutoff; x2++)
	if(y+y2>=0 && y+y2< scaleH && x+x2 >=0 && x+x2 < scaleW && scaleMask[x + x2 + scaleW*(y+y2)] == 1){
		loc = ((y2+pcutoff)*pmapW) + (x2+pcutoff);
		pixels3[pxloc] += pixels2[x+x2 + (scaleW*(y+y2))] * pgausMap[loc];
		counts2[pxloc] += pgausMap[loc];
	}
}


for(int i=0; i<scaleH*scaleW; i++)
if(scaleMask[i] == 1)
	pixels2[i] = pixels3[i]/counts2[i];


//-----------------------------Bicubic interpolation--------------------------------

if(interpolate == true)
for(int y=-1; y<scaleH; y++)
for(int x=-1; x<scaleW; x++){

	if(scaleRad > 1){
		for(int yy = -1; yy<3; yy++)
		for(int xx = -1; xx<3; xx++){
			if(y+yy >= 0
				&& y+yy < scaleH
				&& x+xx >= 0
				&& x+xx < scaleW
				&& scaleMask[(x+xx) + scaleW*(y+yy)] == 1)
				G[xx+1][yy+1] = pixels2[x+xx+scaleW*(y+yy)];
			else G[xx+1][yy+1] = 0.0;
	
			C[xx+1][yy+1] = 0.0;
			T[xx+1][yy+1] = 0.0;
		}
	
		for (int ii = 0 ; ii < 4 ; ii++)    // T = G * MT
		for (int jj = 0 ; jj < 4 ; jj++)
		for (int kk = 0 ; kk < 4 ; kk++)
			T[ii][jj] += G[ii][kk] * M[jj][kk];

		for (int ii = 0 ; ii < 4 ; ii++)    // C = M * T
		for (int jj = 0 ; jj < 4 ; jj++)
		for (int kk = 0 ; kk < 4 ; kk++)
			C[ii][jj] += M[ii][kk] * T[kk][jj];
	} //fi scaleRad >1


	for(int y2 = 0; y2<scaleRad; y2++)
	for(int x2 = 0; x2<scaleRad; x2++)
	if(mask.getPixel(x2+(int) Math.floor(scaleRad/2)+x * scaleRad, y2 + (int) Math.floor(scaleRad/2)+ y*scaleRad) != 0){
		//ix = x2*(0.5/scaleRad); // the scaled XY coordinates within each pixel
		//iy = y2*(0.5/scaleRad);

		if(scaleRad > 1){
			iy = y2/((double)scaleRad); // the scaled XY coordinates within each pixel
			ix = x2/((double)scaleRad);

			td = iy*(iy*(iy*(ix * (ix * (ix * C[0][0] + C[1][0]) + C[2][0]) + C[3][0])	
			                     + (ix * (ix * (ix * C[0][1] + C[1][1]) + C[2][1]) + C[3][1]))
			                  +    (ix * (ix * (ix * C[0][2] + C[1][2]) + C[2][2]) + C[3][2]))
			               +       (ix * (ix * (ix * C[0][3] + C[1][3]) + C[2][3]) + C[3][3]);
			scaleOutPxs[ x2+ (int) Math.floor(scaleRad/2)+x*scaleRad + roiW*(y2+ (int) Math.floor(scaleRad/2)+y*scaleRad)] = (float) td;

		} else { // no interpolation
			td = pixels2[x+scaleW*y];
			scaleOutPxs[ x2+x*scaleRad + roiW*(y2+y*scaleRad)] = (float) td;
		}

		pxSum += td;
		nPx ++;
	}
} else  {//yx mask

for(int y=0; y<scaleH; y++)
for(int x=0; x<scaleW; x++)
if(scaleMask[x + scaleW*y] == 1)
	for(int y2 = 0; y2<scaleRad; y2++)
	for(int x2 = 0; x2<scaleRad; x2++)
	if(mask.getPixel(x2+x*scaleRad, y2 + y*scaleRad) != 0){
		td = pixels2[x+scaleW*y];
		scaleOutPxs[ x2+x*scaleRad + roiW*(y2 +y*scaleRad)] = (float) (td);
		pxSum += td;
		nPx ++;
	}
} // no interpolation


double pxMean = pxSum/nPx;
double pxSSum = 0.0;


for(int y=0; y<roiH; y++)
for(int x=0; x<roiW; x++)
if(mask.getPixel(x, y) != 0){
	pxSSum += Math.pow( scaleOutPxs[x+roiW*y]-pxMean,2);
	pooledPxs[x+roiW*y] += scaleOutPxs[x+roiW*y];
}


//IJ.log("scale:" + k+ " angle:" + j + "mean: " + pxMean + " SD:" + Math.pow(pxSSum/nPx, 0.5));

	// ------------------ OUTPUT RESULTS--------------------------

	ts = ts + "," + Math.pow(pxSSum/nPx, 0.5);

	// if adding new row:
	//rt.incrementCounter();
	//rt.addValue("Title", imTitle);
	//rt.addValue("Scale", k);
	//rt.addValue("Angle", j);
	//rt.addValue("Energy", Math.pow(pxSSum/nPx, 0.5));

	//rt.addValue("s"+k, Math.pow(pxSSum/nPx, 0.5));

outStack.addSlice("SF: " + lumSF[k], scaleOutPxs);

//}//j angles


IJ.showProgress((float) 1);

//new ImagePlus("Scaled Gabor output", scaleOutStack).show();

}//k


//--------------------Print pattern energy (used for setting weights)-----------------------
//IJ.log("Average Energy: " + ts);
IJ.log("Proportion saturated: " + satString);

// output bandpass stack
new ImagePlus("DoG output", outStack).show();

// output pooled image

for(int y=0; y<roiH; y++)
for(int x=0; x<roiW; x++)
if(mask.getPixel(x, y) != 0)
	pooledPxs[x+roiW*y] = (float) (pooledPxs[x+roiW*y]/nOctaves);
else
	pooledPxs[x+roiW*y] = Float.NaN;

ImageStack pooledImage = new ImageStack(roiW, roiH);
pooledImage.addSlice("Pooled Image", pooledPxs);
new ImagePlus("Pooled Image", pooledImage).show();


//rt.showRowNumbers(true);
//rt.show("Results");

}
}
```
