## Supplementary model code for "A model of colour appearance based on efficient coding of natural images": Gabor_ScC_Model_Michelson.java

```
import ij.*;
import ij.process.*;
import ij.gui.*;
import java.awt.*;
import ij.plugin.*;
import ij.util.*;

import ij.measure.ResultsTable;

/*

	Title: Gabor ROI Bandpass
	Author: Jolyon Troscianko
	Date: 17/9/2021

.................................................................................................................

Description:
'''''''''''''''''''''

This script applies a bank of Gabor filters to an ROI. The script has been designed to
work with ROIs of any (non-rectangular) shape by applying a 2D convolution and
weighting all measurements by the degree to which the kernel is inside the ROI
i.e. it re-normalises the kernel with each convolution.

The script efficiently applies the Gabor filter to a series of different spatial scales
by effectively scaling the size of the image (with no interpolation). This makes
it incredibly fast at measuring the larger spatial frequencies that would otherwise
be incredibly slow to process (e.g. by using a larger sigma value).

The script then applies a bicubic interpolation (Catmull-Rom) to the output. This
interpretation assumes that pixels outside the ROI have a value of zero, which
makes sense given the convolution is normalised to have average output of zero.
This is different from a standard image, where out-of-bounds pixels cannot be assumed
to have a particular value.

The result is a very fast convolution, with increasing processing efficiency at larger
spatial scales, combined with smooth output. Any systematic errors introduced by
the scaling and interpolation will be applied equally at all spatial scales.


Input Requirements:
'''''''''''''''''''''
Requires a 32-bit image. If an ROI is selected it will use it, otherwise it will
process the whole image.

The size of the selection must be larger than 2^nOctaves otherwise
the entire image will be a single pixel at larger scales. If the SD ("energy")
is zero at a given scale then avoid using that scale or greater, and probably
avoid using the first smaller scale.


Bicubic interpolation: https://mrl.cs.nyu.edu/~perlin/java/Bicubic.html


*/


public class Gabor_ScC_Model_Michelson implements PlugIn {
public void run(String arg) {

double negM = 1.1; // in a natural scene, unclipped, the negative median is almost exacly 2 times larger than the positive (for all but lowes SFs where sampling error comes in)

double DoGMmichelson = 1.50; // DoG: 2.75 Gabor: around 1.5 to 1.25 is right
double DoGM = 1.50; // approx 0.04 cuts off contrast, 0.06


/*

DoG multiplier was calculated from using images of sinewaves with a frequency gradient
to ellicit the maximum response. Seems to be roughly 1 with relative mordulation.


*/


//-----------------CS Values from Kim et al 2013----------------------
//double[]  lumSF = {0.125,0.25,0.5,1,2,4,8,16};
//double[] lumCS = {21.77,39.98,86.14,197.83,266.64,229.67,181.65,37.91};
// order matching k
//double[] lumCS = {37.91, 181.65, 229.67, 266.64, 197.83, 86.14, 39.98, 21.77};

//weights based on DoGM 1390.3, 19 degree (1730px) woodland, averaged across angles, DR110 SR1.55
//double[] lumWeights = {0.091139756031568,0.076438164921595,0.067983991067059,0.066303333942748,0.078591987268566,0.093507216620015,0.121373933031768,0.198489217699979};
// OLD
//weights based on DoGM 1390.3, 19 degree (1730px) woodland, averaged across angles, DR51 SR1.5
//double[] lumWeights = {0.094627287910463,0.056293963481582,0.047579748697138,0.044395575905736,0.0557900465247,0.080168470269305,0.118314804720173,0.204513065587049};
//weights based on DoGM 2.5, 19 degree (1730px) woodland (linear, *100 +1 log), averaged across angles, DR51 SR1.5
//double[] lumWeights = {0.02858535,0.02604837925,0.02252563875,0.02149476925,0.023615579,0.02363561425,0.01959428,0.02312790725};

//First itteration weights based on DoGM 1.25, 19 degree (1730px) woodland linear, averaged across angles, DR17 SR2.2
//double[] lumWeights = {0.03588935504071984,0.013372979139657258,0.011076214514898534,0.010359056904120536,0.012908562268567784,0.02147841427200531,0.030488495172715617,0.06666620658214883};
//Second itteration of above with updated crispening fit DR19 PS1.5
//double[] lumWeights = {0.06825525926900933,0.025359089066583084,0.020560398175285682,0.01866144278075617,0.024054405277268195,0.03959559748267545,0.05163119044886268,0.09319851130565146};


	ImagePlus imp = IJ.getImage();

	// ---------------------- Calculate Gabor Filter Stuff-------------------------------

	int nAngles = 4;
	double sigma = 2.0;
	double gamma = 1.0;
	double Fx = 3.0;
	//int nOctaves = 6;
	String imTitle = imp.getTitle();
	double dr = 3.75;
	double postSigma = 0.95;
	double poolingWeight = 1.0;

	GenericDialog gd = new GenericDialog("GabRat Disruption Measurement");
		gd.addMessage("Gabor Filter Settings");
		gd.addNumericField("Number_of_angles", nAngles, 0);
		gd.addNumericField("Sigma", sigma, 2);
		gd.addNumericField("Gamma aspect ratio", gamma, 2);
		gd.addNumericField("Frequency", Fx, 2);
		//gd.addNumericField("Number_of_octaves", nOctaves, 0);
		gd.addCheckbox("Michelson", true);
		gd.addCheckbox("Clipping", true);
		gd.addNumericField("Dynamic range", dr, 2);
		gd.addNumericField("postSigma", postSigma, 2);
		gd.addStringField("Spatial_frequency", "0.125,0.25,0.5,1,2,4,8,16", 50);
		// LUM
		gd.addStringField("CSF", "10.885,19.99,43.07,98.915,133.32,114.835,90.825,18.955", 50); // values from kim et al 2016 divided by two, for use with michelson contrast
	
		// RG:
		//gd.addStringField("CSF", "307.3,344.8,307.3,261.5,154.0,97.2,0,0", 50); // Kim et al 2013 values converted from their "C" number to michelson contrast. See spreadsheet for conversion

		gd.addStringField("Gain", "0.0191,0.0181,0.0129,0.0066,0.0052,0.0062,0.0080,0.0232", 50); // Gabor DR2.3 PS1.4 DoGMMich 1.5 Natural scene database
		//gd.addStringField("Gain", "1,1,1,1,1,1,1,1", 50);
		gd.addNumericField("Pooling weight", poolingWeight, 2);
		gd.addCheckbox("Interpolation", true);
		gd.addMessage("Output Images");
		gd.addCheckbox("Gabor Kernel", false);
		gd.addStringField("Label", imTitle, 20);

	gd.showDialog();
	if (gd.wasCanceled())
		return;

	
	nAngles = (int) Math.round(gd.getNextNumber());
	sigma = gd.getNextNumber();
	gamma = gd.getNextNumber();
	Fx = gd.getNextNumber();

	boolean michelson = gd.getNextBoolean();
	boolean clipping = gd.getNextBoolean();

	dr = gd.getNextNumber();
	postSigma = gd.getNextNumber();


// Import input data
	String[] lumSFstring = Tools.split(gd.getNextString(), ",");
	//----------convert string to numeric array-----------
	double[] lumSF = new double[lumSFstring.length];
	double peakSF = 0.0;
	for(int i=0; i<lumSFstring.length; i++){
		lumSF[i] = Double.valueOf(lumSFstring[i]);
		if(lumSF[i] > peakSF)
			peakSF = lumSF[i]; // save peak for use in calculating the scale
	}
	

	String[] lumCSstring = Tools.split(gd.getNextString(), ",");
	//----------convert string to numeric array-----------
	double[] lumCS = new double[lumCSstring.length];
	for(int i=0; i<lumCSstring.length; i++){
		lumCS[i] = Double.valueOf(lumCSstring[i]);
		//IJ.log("SF: " + lumSF[i] + " CS: " + lumCS[i]);
	}

	String[] lumWeightsstring = Tools.split(gd.getNextString(), ",");
	//----------convert string to numeric array-----------
	double[] lumWeights = new double[lumWeightsstring.length];
	for(int i=0; i<lumWeightsstring.length; i++)
		lumWeights[i] = Double.valueOf(lumWeightsstring[i]);
		
	if(lumWeights.length != lumCS.length || lumWeights.length != lumSF.length){
		IJ.log("Error - input value lengths don't match, quitting");
		return;
	}

	int nOctaves = lumCS.length;

	poolingWeight = gd.getNextNumber();
	boolean interpolate = gd.getNextBoolean();
	boolean outputKernel = gd.getNextBoolean();

	imTitle = gd.getNextString();


//---normalise gain/weights array to mean=1------

double meanLumWeight = 0.0;
for(int i=0; i<lumWeights.length; i++)
	meanLumWeight += lumWeights[i];

meanLumWeight = meanLumWeight/lumWeights.length;

for(int i=0; i<lumWeights.length; i++)
	lumWeights[i] = lumWeights[i]/meanLumWeight;

for(int i=0; i<lumWeights.length; i++)
	lumWeights[i] = lumWeights[i]*poolingWeight + (1-poolingWeight);


//-------Kernel stuff-----------
	double psi = Math.PI; // Phase
	if(nAngles/2 != Math.round(nAngles/2))
		nAngles = nAngles -1;

	double sigma_x = sigma;
	double sigma_y = sigma / gamma;
	double largerSigma = 0.0; 

	// Decide size of the filters based on the sigma
	if(sigma_x > sigma_y)
		largerSigma = sigma_x;
	else largerSigma = sigma_y;

	if(largerSigma < 1)
		largerSigma = 1;

	double sigma_x2 = sigma_x * sigma_x;
	double sigma_y2 = sigma_y * sigma_y;

	int filterSizeX = (int) Math.round(6 * largerSigma + 1);
	int filterSizeY = (int) Math.round(6 * largerSigma + 1);
	int filterDim = filterSizeX*filterSizeY;

	double[] kernelArray = new double[(nAngles+1) * filterDim];
	double[] absKernelArray = new double[(nAngles+1) * filterDim];

	int middleX = (int) Math.round(3 * largerSigma);
	int middleY = (int) Math.round(3 * largerSigma);

	//IJ.log("filter x: " + filterSizeX + " y: " + filterSizeY);
	//IJ.log("middle x: " + middleX + " y: " + middleY);


	double rotationAngle = Math.PI/nAngles;
	double theta = 0.0;
	double xPrime = 0.0;
	double yPrime = 0.0;
	double a = 0.0;
	double c = 0.0;

	
	ImageStack kernelStack = new ImageStack(filterSizeX, filterSizeY);
	
	for(int i=0; i<nAngles; i++){
		theta = rotationAngle * i;
		float[] kernelOutputArray = new float[filterSizeX * filterSizeY];

		for(int y=0; y<filterSizeY; y++){
			for(int x=0; x<filterSizeX; x++){
				xPrime = (x-middleX) * Math.cos(theta) + (y-middleY) * Math.sin(theta);
				yPrime = (y-middleY) * Math.cos(theta) - (x-middleX) * Math.sin(theta);
				a = 1.0 / ( 2.0 * Math.PI * sigma_x * sigma_y ) * Math.exp(-0.5 * (xPrime*xPrime / sigma_x2 + yPrime*yPrime / sigma_y2) );
				c = Math.cos( 2.0 * Math.PI * (Fx * xPrime) / filterSizeX + psi);
             				kernelArray[(i*filterDim)+(y*filterSizeX)+x] = -a*c;
             				absKernelArray[(i*filterDim)+(y*filterSizeX)+x] = Math.abs(a*c);
				kernelOutputArray[(y*filterSizeX)+x] = (float) (a*c);
			}//x
		}//y

		if(outputKernel == true)
			kernelStack.addSlice("angle " + i, kernelOutputArray);
		
	}// i (angles)

	if(outputKernel == true)
		new ImagePlus("Gabor Kernel", kernelStack).show();


//----------------calculate size of Gaussian (post-clipping) kernel---------------

double pgVal = (1/Math.pow(2*Math.PI*Math.pow(postSigma,2),0.5)) * Math.exp(-1*(Math.pow(0,2)/(2*Math.pow(postSigma,2) ) ));
double pminVal = 0.001; // this is the cutoff as a proportion of the peak at which it is assumed the Guassian curve is small enough to have no influence
pminVal = pminVal / ( (1/Math.pow(2*Math.PI*Math.pow(postSigma,2),0.5)) * Math.exp(-1*(Math.pow(0,2)/(2*Math.pow(postSigma,2) ) ))); // minVal as a proportaion of start val
int pcutoff = 1;

while(pgVal > pminVal){
	pgVal = (1/Math.pow(2*Math.PI*Math.pow(postSigma,2),0.5)) * Math.exp(-1*(Math.pow(pcutoff,2)/(2*Math.pow(postSigma,2) ) ));
	pcutoff++;
}

// Gaussian Kernel
int pmapW = 2*pcutoff +1;
double[] pgausMap = new double[pmapW*pmapW]; 
double prad = 0.0;

for(int y=0; y<pmapW; y++)
for(int x=0; x<pmapW; x++){
	prad = Math.pow( Math.pow(x-pcutoff,2) + Math.pow(y-pcutoff,2) ,0.5);
	pgausMap[(y*pmapW)+x] = (1/Math.pow(2*Math.PI*Math.pow(postSigma,2),0.5)) * Math.exp(-1*(Math.pow(prad,2)/(2*Math.pow(postSigma,2) ) ));
}


// --------------------- Get ROI Mask---------------------------------


	Roi roi = imp.getRoi();
	if (roi!=null && !roi.isArea()) roi = null;
	ImageProcessor ip = imp.getProcessor();
	int w = ip.getWidth();
	int h = ip.getHeight();
	int roiW = 0;
	int roiH = 0;
	int roiX = 0;
	int roiY = 0;

	//ImageProcessor mask = new ByteProcessor(roiW, roiH);

	if (roi==null) // in case there is no ROI selected use the entire image
		roi = new Roi(0, 0, w, h);
	
	Rectangle r = roi.getBounds();
	ImageProcessor mask = roi.getMask();

	//int roiType = roi.getType();
	//if (roiType==Roi.RECTANGLE){
	if(mask==null){
		//IJ.log("rectangle");
		mask = new ByteProcessor(r.width, r.height);
		mask.invert();
	}

	roiW = r.width;
	roiH = r.height;
	roiX = r.x;
	roiY = r.y;


ImageStack outStack = new ImageStack(roiW, roiH);
double kernelSum = 0.0; // the kernel needs re-normalising with each pass
int kernelCount = 0;
double absKernelSum = 0.0;
double convSum = 0.0;
double absConvSum = 0.0;
double rawSum = 0.0;
double tpx = 0.0;


float[] pixels = new float[roiW*roiH];

for (int y=0; y<roiH; y++)
for (int x=0; x<roiW; x++)
if(mask.getPixel(x,y) != 0){
	pixels[x+ y*roiW] = ip.getPixelValue(x+roiX, y+roiY);
}


//ResultsTable rt =  ResultsTable.getResultsTable();
//rt.incrementCounter();
//rt.addValue("Title", imTitle);


//---------------Bicubic interpolation coefficients----------------
double[][] M = {             // Catmull-Rom basis matrix
	{-0.5,  1.5, -1.5, 0.5},
	{ 1  , -2.5,  2  ,-0.5},
	{-0.5,  0  ,  0.5, 0  },
	{ 0  ,  1  ,  0  , 0  }
};

double[][] C = new double[4][4];
double[][] T = new double[4][4];
double[][] G = new double[4][4];

double ix = 0.0;
double iy = 0.0;


//-------------------------------- downscale pixels-------------------------------------
// sampling at an octave scale

float[] pooledPxs = new float[roiW*roiH];

double aTpos = 0.0;
double aTneg = 0.0;
double sTpos = 0.0;
double sTneg = 0.0;

int scaleRad = 1;
String ts = "";
String tss = "";
double aveEnergy = 0.0;

for(int k=0; k<nOctaves; k++)
if(lumCS[k] > 0.0){

//scaleRad = (int) Math.pow(2, k);
scaleRad = (int) Math.round(peakSF / lumSF[k]);

int scaleW = (int) (Math.floor(roiW/scaleRad) +1);
int scaleH = (int) (Math.floor(roiH/scaleRad) +1);

double scaleSum = 0.0;
int scaleCount = 0;


float[] scalePixels = new float[scaleW*scaleH];
int[] scaleMask = new int[(scaleW)*(scaleH)];

for(int y=0; y<scaleH; y++)
for(int x=0; x<scaleW; x++){

	scaleSum = 0.0;
	scaleCount = 0;

	for(int y2 = 0; y2<scaleRad; y2++)
	for(int x2 = 0; x2<scaleRad; x2++)
	if(mask.getPixel(x2+x*scaleRad,y2+y*scaleRad) != 0){
		scaleSum += pixels[(x2+x*scaleRad) + (roiW*(y2+y*scaleRad))];
		scaleCount ++;
	} //x2 y2 fi

	if(scaleCount > 0){
		scalePixels[x+y*scaleW] = (float) (scaleSum/scaleCount);
		scaleMask[x+y*scaleW] = 1;
	}
} //xy


//-----------------Scale convolution-------------

//ImageStack scaleOutStack = new ImageStack(scaleW, scaleH);

for(int j=0; j<nAngles; j++){


double[] pixels2 = new double[scaleW*scaleH];
int[] counts = new int[scaleW*scaleH];
float[] outPxs = new float[scaleW*scaleH];

int loc = 0;
int pxloc = 0;
int scaleLoc = 0;

for(int y=0; y<scaleH; y++)
for(int x=0; x<scaleW; x++)
if(scaleMask[x + scaleW*y] == 1){

	pxloc = (y*scaleW)+x;

	rawSum = 0.0;
	convSum = 0.0;
	absConvSum = 0.0;
	kernelSum = 0.0;
	absKernelSum = 0.0;
	kernelCount = 0;

	for(int y2=-middleY; y2<=middleY; y2++)  
	for(int x2=-middleX; x2<=middleX; x2++){
		if(y+y2>=0 && y+y2< scaleH && x+x2 >=0 &&x+x2 < scaleW && scaleMask[x + x2 + scaleW*(y+y2)] == 1){
			loc = (j*filterDim) + ((y2+middleY)*filterSizeX) + (x2+middleX);

			tpx = scalePixels[x+x2 + (scaleW*(y+y2))];
			rawSum += tpx;

			kernelSum += kernelArray[loc];
			kernelCount++;
			convSum += tpx * kernelArray[loc];

			if(michelson == true){
				absKernelSum += absKernelArray[loc];
				absConvSum  += tpx * absKernelArray[loc];
			}
		}
	}

	//pixels2[pxloc] = DoGMultiplier * ((convSum-(kernelSum*(rawSum/kernelCount)))  / (absConvSum/2));

	if(michelson == true)
		pixels2[pxloc] = DoGMmichelson * ((convSum-(kernelSum*(rawSum/kernelCount)))  / absConvSum); // Michelson contrast
	else
		pixels2[pxloc] = DoGM * (convSum-(kernelSum*(rawSum/kernelCount)));

	IJ.showProgress((float) y/roiH );
	
} // xy fi


//for(int i=0; i<scaleH*scaleW; i++)
//	outPxs[i] = (float) (pixels2[i]/counts[i]);

//scaleOutStack.addSlice("output angle" + j, outPxs);


//-----------------Apply to original scale (allows for easier measurement and weighting----------


float[] scaleOutPxs = new float[roiW*roiH];

for(int i=0; i<roiW*roiH; i++)
	scaleOutPxs[i] = Float.NaN;

double td = 0.0;
double pxSum = 0.0;
double nPx = 0.0;

//for(int i=0; i<scaleW*scaleH; i++)
//	pixels2[i] = DoGMultiplier*pixels2[i]/counts[i];


// -------------------activation and saturation thresholding---------------

if(clipping == true){

//aTpos = (1/lumCS[k]);
//aTneg = aTpos*-1.0;
//sTpos = aTpos*dr; // saturation threshold
//sTneg = sTpos*-1.0;

aTpos = (1/lumCS[k]);
aTneg = aTpos*-negM;
sTpos = aTpos*dr; // saturation threshold
sTneg = aTneg*dr;

for(int i=0; i<scaleW*scaleH; i++){

		td = pixels2[i];

		double tvv = 0.0;
		
		if(td > aTpos)
			tvv = td - aTpos;
		else if(td > 0)
			tvv = 0.0;
		else if(td < aTneg)
			tvv = td - aTneg;
		else if(td < 0)
			tvv = 0.0;
		else tvv = 0.0;

		if(tvv > sTpos)
			tvv = sTpos;
		if(tvv < sTneg)
			tvv = sTneg;

		pixels2[i] = tvv/lumWeights[k];

}//i


}// clipping ture


//-------------------------------------------------------------------------------------------
//--------------------------Post-clipping Gaussian smoothing--------------------------------


double[] pixels3 = new double[scaleW*scaleH];
double[] counts2 = new double[scaleW*scaleH];

for(int y=0; y<scaleH; y++)
for(int x=0; x<scaleW; x++){
	pxloc = (y*scaleW)+x;
	if(scaleMask[x + scaleW*y] == 1)
	for(int y2=-pcutoff; y2<=pcutoff; y2++)  
	for(int x2=-pcutoff; x2<=pcutoff; x2++)
	if(y+y2>=0 && y+y2< scaleH && x+x2 >=0 && x+x2 < scaleW && scaleMask[x + x2 + scaleW*(y+y2)] == 1){
		loc = ((y2+pcutoff)*pmapW) + (x2+pcutoff);
		pixels3[pxloc] += pixels2[x+x2 + (scaleW*(y+y2))] * pgausMap[loc];
		counts2[pxloc] += pgausMap[loc];
	}
}


for(int i=0; i<scaleH*scaleW; i++)
if(scaleMask[i] == 1)
	pixels2[i] = pixels3[i]/counts2[i];


//-----------------------------Bicubic interpolation--------------------------------

if(interpolate == true)
for(int y=-1; y<scaleH; y++)
for(int x=-1; x<scaleW; x++){

	if(scaleRad > 1){
	//if(scaleRad > 1 && interpolate == true){
		for(int yy = -1; yy<3; yy++)
		for(int xx = -1; xx<3; xx++){
			if(y+yy >= 0
				&& y+yy < scaleH
				&& x+xx >= 0
				&& x+xx < scaleW
				&& scaleMask[(x+xx) + scaleW*(y+yy)] == 1)
				G[xx+1][yy+1] = pixels2[x+xx+scaleW*(y+yy)];
			else G[xx+1][yy+1] = 0.0;
	
			C[xx+1][yy+1] = 0.0;
			T[xx+1][yy+1] = 0.0;
		}
	
		for (int ii = 0 ; ii < 4 ; ii++)    // T = G * MT
		for (int jj = 0 ; jj < 4 ; jj++)
		for (int kk = 0 ; kk < 4 ; kk++)
			T[ii][jj] += G[ii][kk] * M[jj][kk];

		for (int ii = 0 ; ii < 4 ; ii++)    // C = M * T
		for (int jj = 0 ; jj < 4 ; jj++)
		for (int kk = 0 ; kk < 4 ; kk++)
			C[ii][jj] += M[ii][kk] * T[kk][jj];
	} //fi scaleRad >1

	for(int y2 = 0; y2<scaleRad; y2++)
	for(int x2 = 0; x2<scaleRad; x2++)
	if(mask.getPixel(x2+(int) Math.floor(scaleRad/2)+x * scaleRad, y2 + (int) Math.floor(scaleRad/2)+ y*scaleRad) != 0){
		//ix = x2*(0.5/scaleRad); // the scaled XY coordinates within each pixel
		//iy = y2*(0.5/scaleRad);

		//if(scaleRad > 1 && interpolate == true){
		if(scaleRad > 1){
			iy = y2/((double)scaleRad); // the scaled XY coordinates within each pixel
			ix = x2/((double)scaleRad);

			td = iy*(iy*(iy*(ix * (ix * (ix * C[0][0] + C[1][0]) + C[2][0]) + C[3][0])	
			                     + (ix * (ix * (ix * C[0][1] + C[1][1]) + C[2][1]) + C[3][1]))
			                  +    (ix * (ix * (ix * C[0][2] + C[1][2]) + C[2][2]) + C[3][2]))
			               +       (ix * (ix * (ix * C[0][3] + C[1][3]) + C[2][3]) + C[3][3]);
			//scaleOutPxs[ x2+ (int) Math.floor(scaleRad/2)+x*scaleRad + roiW*(y2+ (int) Math.floor(scaleRad/2)+y*scaleRad)] = (float) td;
			scaleOutPxs[ x2+ (int) Math.floor(scaleRad/2)+x*scaleRad + roiW*(y2+ (int) Math.floor(scaleRad/2)+y*scaleRad)] = (float) td;

		//} else if(interpolate == false){
		//	td = pixels2[x+scaleW*y];
		//	scaleOutPxs[ x2+ (int) Math.floor(scaleRad/2)+x*scaleRad + roiW*(y2+ (int) Math.floor(scaleRad/2)+y*scaleRad)] = (float) td;
		}else { // no interpolation
			td = pixels2[x+scaleW*y];
			scaleOutPxs[ x2+x*scaleRad + roiW*(y2+y*scaleRad)] = (float) td;
		}

		pxSum += td;
		nPx ++;
	}

} else  {//yx mask

for(int y=0; y<scaleH; y++)
for(int x=0; x<scaleW; x++)
if(scaleMask[x + scaleW*y] == 1)
	for(int y2 = 0; y2<scaleRad; y2++)
	for(int x2 = 0; x2<scaleRad; x2++)
	if(mask.getPixel(x2+x*scaleRad, y2 + y*scaleRad) != 0){
		td = pixels2[x+scaleW*y];
		scaleOutPxs[ x2+x*scaleRad + roiW*(y2 +y*scaleRad)] = (float) (td);
		pxSum += td;
		nPx ++;
	}
} // no interpolation


double pxMean = pxSum/nPx;
double pxSSum = 0.0;


for(int y=0; y<roiH; y++)
for(int x=0; x<roiW; x++)
if(mask.getPixel(x, y) != 0){
	pxSSum += Math.pow( scaleOutPxs[x+roiW*y]-pxMean,2);
	pooledPxs[x+roiW*y] += scaleOutPxs[x+roiW*y];
}


//IJ.log("scale:" + k+ " angle:" + j + "mean: " + pxMean + " SD:" + Math.pow(pxSSum/nPx, 0.5));

	// ------------------ OUTPUT RESULTS--------------------------

	aveEnergy += Math.pow(pxSSum/nPx, 0.5);
	ts = ts + "," + Math.pow(pxSSum/nPx, 0.5);


	// if adding new row:
	//rt.incrementCounter();
	//rt.addValue("Title", imTitle);
	//rt.addValue("Scale", k);
	//rt.addValue("Angle", j);
	//rt.addValue("Energy", Math.pow(pxSSum/nPx, 0.5));

	//rt.addValue("s"+k+"_a"+j, Math.pow(pxSSum/nPx, 0.5));

outStack.addSlice("scale" +k + "_angle" + j, scaleOutPxs);

}//j angles

tss = tss + "," + (aveEnergy/nAngles);
aveEnergy = 0.0;


//IJ.log("LumCS: " + lumCS[k] + "aT: " + aTpos + " sT: " + sTpos);

IJ.showProgress((float) 1);

//new ImagePlus("Scaled Gabor output", scaleOutStack).show();

}//k


//--------------------Print pattern energy (used for setting weights)-----------------------
//IJ.log("Oriented Energy: " + ts);
//IJ.log("Average Energy: " + tss);

new ImagePlus("Gabor output", outStack).show();

// output pooled image

for(int y=0; y<roiH; y++)
for(int x=0; x<roiW; x++)
if(mask.getPixel(x, y) != 0)
	pooledPxs[x+roiW*y] = (float) (pooledPxs[x+roiW*y]/nOctaves);
else
	pooledPxs[x+roiW*y] = Float.NaN;

ImageStack pooledImage = new ImageStack(roiW, roiH);
pooledImage.addSlice("Pooled Image", pooledPxs);
new ImagePlus("Pooled Image", pooledImage).show();


//rt.showRowNumbers(true);
//rt.show("Results");

}
}
```
