## Supplementary model code for "A model of colour appearance based on efficient coding of natural images": Instructions.txt

The Spatiochromatic Bandwidth Limited model was developed by Jolyon Troscianko and Daniel Osorio

This code is provided without warrany - use at your own risk.

To install and use the code:

1) Install ImageJ from NIH (version 1.52+)

2) Copy these files into a directory within your ImageJ/plugins/ directory, e.g. ImageJ/plugins/SBL/

3) Open an image (e.g. conventional sRGB, 24-bit image) and go "plugins>SBL>SBL Model Gabor" to run the model. You should ensure the input image is of an appropriate scale for your modelled viewing distance. e.g. for the DoG model the peak wavelength sensitivity is 5.7 pixels, and the highest SF in the model is 16 cycles per degree. So the angular width of the image can be scaled to this. e.g. the Whittle screen was 19 degrees wide, which is 19 degrees * 16 cycles per degree=304 cycles, 5.7*304=1733, so the image should be scaled to this width (i.e. 1733 pixels wide is 19 degrees).

Further info: The "SBL_Model_DoG.ijm" and "SBL_Model_Gabor.ijm" scripts can be modified to process specific images. The SBL model itself is run by the "DoG_ScC_Model_Michelson.java" and "Gabor_ScC_Model_Michelson.java" scripts.
