## Supplementary model code for "A model of colour appearance based on efficient coding of natural images": Michelson_opponent_B.java

```
/*__________________________________________________________________

	Title: 
	''''''''''''''''''''''''''''''''''''''''''''''''''''''''''''''''''''''''''''''''''''''''
	Author: Jolyon Troscianko
	Date: 
............................................................................................................


____________________________________________________________________
*/


import ij.*;
import ij.plugin.filter.PlugInFilter;
import ij.process.*;

public class Michelson_opponent_B implements PlugInFilter {

ImageStack stack;
	public int setup(String arg, ImagePlus imp) { 
	stack = imp.getStack(); 
	return DOES_32 + STACK_REQUIRED; 
	}
public void run(ImageProcessor ip) {

float[] R;
float[] G;
float[] B;

int w = stack.getWidth();
int h = stack.getHeight();
int dimension = w*h;

R = (float[]) stack.getPixels(1);
G = (float[]) stack.getPixels(2);
B = (float[]) stack.getPixels(3);

float[] Lum = new float[dimension];
float[] RG = new float[dimension];
float[] YB = new float[dimension];

double Y = 0.0;
double rg = 0.0;
double yb = 0.0;


for(int i=0; i<dimension; i++){


	Y = (R[i]+G[i])/2;
	Lum[i] = (float) (R[i]*0.629 + G[i]*0.314 + B[i]*0.057); // ratios from Hofer et al 2005
	//Lum[i] = (float) Y;
	if(R[i] == 0 && G[i] == 0)
		rg = (float) 0.0;
	else{
		rg = ((R[i]-G[i])/(R[i]+G[i]));
	//	if(rg > 1.0)
	//		rg=1.0;
	//	if(rg<-1.0)
	//		rg=-1.0;
	}


	if(Y == 0 && B[i] == 0)
		YB[i] = (float) 0.0;
	else{
		yb = (float) ((Y-B[i])/(Y+B[i]));
	//	if(yb > 1.0)
	//		yb = 1.0;
	//	if(yb<-1.0)
	//		yb = -1.0;
	}

	RG[i] = (float) rg;
	YB[i] = (float) yb;
	
}//i


ImageStack outStack = new ImageStack(w, h);
outStack.addSlice("Lum", Lum);
outStack.addSlice("RG", RG);
outStack.addSlice("YB", YB);
new ImagePlus("Michelson Opponent Image", outStack).show();


}
}
```
