## Supplementary model code for "A model of colour appearance based on efficient coding of natural images": MichelsonContrast_to_Cone_Catch.java

```
/*__________________________________________________________________

	Title: 
	''''''''''''''''''''''''''''''''''''''''''''''''''''''''''''''''''''''''''''''''''''''''
	Author: Jolyon Troscianko
	Date: 
............................................................................................................


Y = (R+G)/2
lum = ln(Y)          (should really be average weighted by all cone ratios)
RG = ln((R-G)/(R+G)) = ln(R-G)-ln(R+G)
BY = ln((Y-B)/(Y+B)) = ln(Y-B)-ln(Y+B)

____________________________________________________________________
*/


import ij.*;
import ij.plugin.filter.PlugInFilter;
import ij.process.*;

public class MichelsonContrast_to_Cone_Catch implements PlugInFilter {

ImageStack stack;
	public int setup(String arg, ImagePlus imp) { 
	stack = imp.getStack(); 
	return DOES_32 + STACK_REQUIRED; 
	}
public void run(ImageProcessor ip) {

float[] Lum;
float[] RG;
float[] YB;

int w = stack.getWidth();
int h = stack.getHeight();
int dimension = w*h;

Lum = (float[]) stack.getPixels(1);
RG = (float[]) stack.getPixels(2);
YB = (float[]) stack.getPixels(3);

float[] R = new float[dimension];
float[] G = new float[dimension];
float[] B = new float[dimension];

double Y = 0.0;
double r = 0.0;
double g = 0.0;
double b = 0.0;
double expVal = 1;


for(int i=0; i<dimension; i++){

/*
	Y = Math.exp(Lum[i]);
	B[i] = (float) (Y / Math.exp(YB[i]) );
	R[i] = (float) ( Y+Y*((Math.exp(RG[i])-1) / (Math.exp(RG[i])+1))   );
	G[i] = (float) ( R[i] / Math.exp(RG[i]) );
*/

	r =(Lum[i]*2)/(1+ (1-RG[i])/(1+RG[i]) );
	//R[i] = (float) Math.pow(r, expVal);
	//r =(Lum[i]*2)/(1+ (1-Math.exp(RG[i]))/(1+Math.exp(RG[i])) );
	R[i] = (float) r;

	g = r/((1+RG[i])/(1-RG[i]));
	//G[i] = (float) Math.pow(g, expVal);
	//g = r/((1+Math.exp(RG[i]))/(1-Math.exp(RG[i])));
	G[i] = (float) g;
	
	b = (Lum[i]*(1+(1-YB[i])/(1+YB[i])))-Lum[i];
	//B[i] = (float) Math.pow(b, expVal);
	//b = (Math.exp(Lum[i])*(1+(1-Math.exp(YB[i]))/(1+Math.exp(YB[i]))))-Math.exp(Lum[i]);
	B[i] = (float) b;
	
}//i


ImageStack outStack = new ImageStack(w, h);
outStack.addSlice("R", R);
outStack.addSlice("G", G);
outStack.addSlice("B", B);
new ImagePlus("ConeCatch Image", outStack).show();


}
}
```
