## Supplementary model code for "A model of colour appearance based on efficient coding of natural images": SBL_Model_DoG.ijm

```
lumOnly = 0;
showBandpass = 0;
poolW = 1;
postSigma = 0.95;

// ------------ CSFs---------------
SFs ="0.125,0.25,0.5,1,2,4,8,16";
lumCS = "10.885,19.99,43.07,98.915,133.32,114.835,90.825,18.955"; // kim et al 2013 divided by two (from relative modulation michelson)
rgCS = "326.34,326.34,391.83,370.11,269.88,187.68,92.72,0"; // Kim et al 2013 values converted from their "C" number to michelson contrast. See spreadsheet for conversion. Based on average data from observers that tested the entire SF range
//rgCS = "114.218,114.218,137.141,129.540,94.458,65.686,32.450,0";
byCS = "32.76,32.76,40.18,42.20,45.34,38.69,14.00,0";


//------------GAIN---------------
//lumGain = "0.205,0.244,0.261,0.181,0.147,0.170,0.206,0.367"; // based on crispening fit DR20
//lumGain = "0.0191,0.0181,0.0129,0.0066,0.0052,0.0062,0.0080,0.0232"; // Gabor DR2.3 PS1.4 DoGMMich 1.5 Natural scene database
lumGain = "0.201,0.221,0.179,0.097,0.074,0.087,0.109,0.309";

//rgGain = "1,1,1,1,1,1,1,1";
// using lum for RG
//rgGain = "0.0191,0.0181,0.0129,0.0066,0.0052,0.0062,0.0080,0.0232"; // Using LUM Gabor DR2.3 PS1.4 DoGMMich 1.5 Natural scene database
rgGain = "0.022,0.023,0.021,0.023,0.030,0.038,0.049,0"; // DR15 PS0.75 natural scene weights negM1


// using lum for by
//byGain = "0.0191,0.0181,0.0129,0.0066,0.0052,0.0062,0.0080,0.0232"; // Using LUM Gabor DR2.3 PS1.4 DoGMMich 1.5 Natural scene database
//byGain = "1,1,1,1,1,1,1,1";
byGain = "0.030,0.035,0.041,0.046,0.048,0.054,0.065,0"; // DR5 PS0.75 natural scene weights  negM1


//----------Dynamic Range--------------
lumDR = 15; //15 matches crispening
rgDR = 15;
byDR = 5; 


//--------------Start Image processing---------------------
setBatchMode(true);


if(bitDepth() == 8)
	lumOnly = 1;

if(nSlices < 3 && nSlices > 1)
	lumOnly = 1;

if(bitDepth() == 24){
	//run("sRGB to CIEXYZ");
	run("sRGB to linRGB");
	//run("RGB Stack");
	//run("32-bit");
	//run("Add...", "value=1 stack"); // ensure no zeros
}

if(lumOnly == 0)
	//////////////run("Relative Modulation Opponency");
	//run("Michelson opponent");
	run("Michelson opponent B");

rmID = getImageID(); // relative modulation ID

//w=getWidth();
//h=getHeight();

//LUM
//run("DoG ScC Model Relative Modulation input", "sigma1=1 sigma2=1.60 dynamic=&lumDR post=1.75 spatial_frequency=[&SFs] csf=[&lumCS] gain=[&lumGain] interpolation label=[]");

// run DoG with michelson contrast
run("DoG ScC Model Michelson", "sigma1=1 sigma2=1.60 michelson clipping dynamic=&lumDR post=&postSigma spatial_frequency=[&SFs] csf=[&lumCS] gain=[&lumGain] pooling=&poolW interpolation label=[]");

//run Gabor
//run("Gabor ScC Model Michelson", "number_of_angles=4 sigma=2 gamma=1 frequency=3 michelson clipping dynamic=&lumDR postsigma=&postSigma spatial_frequency=[&SFs] csf=[&lumCS] gain=[&lumGain] pooling=&poolW interpolation label=[]");


if(lumOnly == 1){
	setBatchMode("show");
	exit();
}

lumID = getImageID();


if(lumOnly == 0){
	// LUM
	run("Select All");
	run("Copy");
	close();
	rename("Lum bandpass");
	if(showBandpass == 1)
		setBatchMode("show");
	else close();
	selectImage(rmID);
	run("Paste");
	
	// RG
	setSlice(2);
	//run("DoG ScC Model Relative Modulation input", "sigma1=1 sigma2=1.60 dynamic=&rgDR post=1.75 spatial_frequency=[&SFs] csf=[&rgCS] gain=[&rgGain] interpolation label=[]");
	// run without michelson contrast
	run("DoG ScC Model Michelson", "sigma1=1 sigma2=1.60 clipping dynamic=&rgDR post=&postSigma spatial_frequency=[&SFs] csf=[&rgCS] gain=[&rgGain] pooling=&poolW interpolation label=[]");

	run("Select All");
	run("Copy");
	close();
	rename("RG bandpass");
	if(showBandpass == 1)
		setBatchMode("show");
	else close();
	selectImage(rmID);
	run("Paste");

	//BY
	setSlice(3);
	//run("DoG ScC Model Relative Modulation input", "sigma1=1 sigma2=1.60 dynamic=&byDR post=1.75 spatial_frequency=[&SFs] csf=[&byCS] gain=[&byGain] interpolation label=[]");
	// run without michelson contrast
	run("DoG ScC Model Michelson", "sigma1=1 sigma2=1.60 clipping dynamic=&byDR post=&postSigma spatial_frequency=[&SFs] csf=[&byCS] gain=[&byGain] pooling=&poolW interpolation label=[]");

	run("Select All");
	run("Copy");
	close();
	rename("BY bandpass");
	if(showBandpass == 1)
		setBatchMode("show");
	else close();
	selectImage(rmID);
	run("Paste");
	setSlice(1);
	setBatchMode("show");
	
	// convert back to cone-catch
	//run("Relative Modulation Opponency to Cone Catch");
	//run("CIEXYZ to Linear sRGB");


	run("Duplicate...", "duplicate");
	rename("temp");

	setSlice(1);
	run("Add...", "value=1 slice");

	setSlice(2);
	run("Multiply...", "value=2 slice"); //0.25
	setSlice(3);
	run("Multiply...", "value=2 slice"); //0.25

	run("MichelsonContrast to Cone Catch");


	powVal = 0.8;
	ts = "code=v=pow(v," + powVal + ")";

	setSlice(1);
	run("Macro...", ts);
	setSlice(2);
	run("Macro...", ts);
	setSlice(3);
	run("Macro...", ts);

	run("Make Composite", "display=Composite");
	setSlice(1);
	setMinAndMax("0.75", "1.06");
	setSlice(2);
	setMinAndMax("0.75", "1.06");
	setSlice(3);
	setMinAndMax("0.75", "1.06");
	setBatchMode("show");
}

// exp lum plus 1, also divid RG by 4 to get similar SD? then make non-linear again, e.g. pow0.4
```
