## Supplementary model code for "A model of colour appearance based on efficient coding of natural images": sRGB_to_linRGB.java

```
/*__________________________________________________________________

	Title: sRGB to linearRGB
	''''''''''''''''''''''''''''''''''''''''''''''''''''''''''''''''''''''''''''''''''''''''
	Author: Jolyon Troscianko
	Date: 21/01/2020
............................................................................................................

Based on the reverse matrix here: https://www.image-engineering.de/library/technotes/958-how-to-convert-between-srgb-and-ciexyz

Note that these are the linear (R'G'B') values and need to go through gamma correction before being true sRGB values

____________________________________________________________________
*/


import ij.*;
import ij.plugin.filter.PlugInFilter;
import ij.process.*;

public class sRGB_to_linRGB implements PlugInFilter {

ImageStack stack;
	public int setup(String arg, ImagePlus imp) { 
	stack = imp.getStack(); 
	return  DOES_RGB; 
	}
public void run(ImageProcessor ip) {


int w = stack.getWidth();
int h = stack.getHeight();
int dimension = w*h;

float[] r = new float[dimension];
float[] g = new float[dimension];
float[] b = new float[dimension];

int[] px =(int[]) stack.getPixels(1);

double R = 0.0;
double G = 0.0;
double B = 0.0;

for (int i=0;i<dimension;i++) {

	R = (double) ((px[i]>>16)&0xff)/255.0;
	G = (double) ((px[i]>>8)&0xff)/255.0;
	B = (double) (px[i]&0xff)/255.0;

//linearise
	if(R <= 0.04045)
		R = R/12.92;
	else R = Math.pow( (R + 0.055)/1.055, 2.4);

	if(G <= 0.04045)
		G = G/12.92;
	else G = Math.pow( (G + 0.055)/1.055, 2.4);

	if(B <= 0.04045)
		B = B/12.92;
	else B = Math.pow( (B + 0.055)/1.055, 2.4);

	r[i] = (float) R;
	g[i] = (float) G;
	b[i] = (float) B;
}


ImageStack outStack = new ImageStack(w, h);
outStack.addSlice("R'", r);
outStack.addSlice("G'", g);
outStack.addSlice("B'", b);
new ImagePlus("Linear RGB", outStack).show();

}
}
```
